# Dissection of a non-coding risk locus at 1p36.23 identifies *ERRFI1* as a novel gene in the pathogenesis of psoriasis and psoriatic arthritis

**DOI:** 10.1101/2023.12.04.569945

**Authors:** Oliver J. Gough, Shraddha S. Rane, Amy Saunders, Megan Priestley, Helen Ray-Jones, Chenfu Shi, Richard B. Warren, Antony Adamson, Stephen Eyre

## Abstract

**Background:** Psoriasis and its associated inflammatory arthritis Psoriatic Arthritis (PsA) are potentially life-ruining conditions associated with numerous comorbidities. A previously-identified genetic risk association for psoriasis and PsA lies in a non-coding region at chromosome 1p36.23, and as such functional validation is required to determine the genetic mechanism contributing to psoriatic disease risk.

**Results:** rs11121131 – a variant in tight linkage with rs11121129, the lead GWAS variant for the 1p36.23 association – lies in a putative enhancer active in keratinocytes but not in immune cells. Promoter-capture Hi-C and H3K27Ac HiChIP showed keratinocyte-specific interactions between 1p36.23 and the *TNFRSF9/PARK7/ERRFI1* gene locus ∼200Kb upstream of the risk locus. Deletion of the enhancer in HaCat keratinocytes led to a reduction in transcript levels of the gene *ERRFI1*, a negative regulator of Epidermal Growth Factor Receptor (EGFR) signalling. CRISPR activation of the enhancer also affected *ERRFI1* levels, but paradoxically showed that steady-state activation led to repression of *ERRFI1*, accompanied by significant deposition of H3K27Me3 histone marks at both the enhancer and the *ERRFI1* gene locus. ERRFI1 levels were shown to be increased in inflamed skin from a mouse model of psoriasis, further suggesting its involvement in disease.

**Conclusions:** These data indicate rs11121131 lies in an enhancer which modulates *ERRFI1* expression in keratinocytes, providing a likely risk mechanism for the 1p36.23 risk association. *ERRFI1* represents a novel gene in the pathogenesis of psoriasis and PsA – improving our understanding of these diseases – and the ERRFI1/EGFR signalling axis may therefore be a target for new treatment modalities for psoriatic disease.

## Background

Genome Wide Association Studies (GWASs) in autoimmune diseases have shown that approximately 90% of genetic variants which contribute to disease risk lie in non-coding regulatory regions of the genome (1, 2). Determining the mechanisms and target genes of these regulatory risk loci is inherently difficult compared to coding variants. Non-coding variants often affect the regulation of distal genes, and can do so by disruption of a variety of regulatory mechanisms (3). Furthermore, due to linkage disequilibrium (LD) between the lead variant and other common variants at GWAS loci, determining the true causal variant for an association remains challenging.

Psoriasis (Ps) is a chronic inflammatory skin condition, with an estimated prevalence of between 0.09% and 5.1% amongst populations worldwide (4). Approximately 1 in 5 psoriasis patients go on to develop an associated inflammatory arthropathy, termed Psoriatic Arthritis (PsA), which is functionally distinct from other common arthritides such as Rheumatoid Arthritis (RA) and Osteoarthritis (OA) (5-7). These conditions can significantly affect quality of life and mortality, and are associated with several comorbidities such as Inflammatory Bowel Disease (IBD) and metabolic syndrome (8-12).

While GWAS and rare variant analyses have identified around 126 loci associated with psoriasis and around 20 directly associated with PsA (13-17), the functional mechanisms underlying many of these risk associations are still not fully understood. This is particularly true for PsA, where clinical heterogeneity and small cohort sizes have limited the power of GWAS (18), and direct functional follow-up of genetic risk loci has been rare. While there are effective treatment strategies for psoriasis, these generally entail systemic immunosuppression – leaving patients at risk of systemic infections (19, 20). For PsA, response to conventional first-line treatments is poor compared to other arthritides (21), and therefore there is an unmet need for new and effective therapies. The clinical heterogeneity of PsA also makes diagnosis and early prediction of onset difficult (22). It is likely that a greater understanding of the functional and genetic basis of these diseases will improve their treatment and diagnosis, and perhaps lead to new treatment modalities.

Advances in CRISPR-Cas technologies have revolutionised the analysis of non-coding risk loci, allowing for direct perturbation of DNA regions in their endogenous genomic context, to empirically validate variant-gene associations (23-25). Here, we apply Hi-C and CRISPR-Cas technologies to the analysis of a psoriasis and PsA risk association at 1p36.23, tagged by the single nucleotide polymorphism (SNP) rs11121129. This was identified via GWAS as a psoriasis risk locus (26, 27), and later replicated in a PsA cohort (28) in studies utilising the Immunochip fine-mapping genotyping array (29). Through our analysis, we find that a SNP in tight LD with the lead GWAS variant lies in a putative keratinocyte-specific enhancer controlling expression of *ERRFI1* – a feedback inhibitor of the Epidermal Growth Factor Receptor (EGFR). Furthermore, we show that there is an increase in levels of ERRFI1 in inflamed skin from a mouse model of psoriasis, providing further evidence for the role of ERRFI1 in psoriatic inflammation. In doing so, this work functionally implicates *ERRFI1* – and its aberrant control in skin cells – as a potential novel gene in the pathogenesis of both Ps and PsA, providing a likely causal gene and variant for the 1p36.23 risk association for these conditions. By implicating *ERRFI1*, this study provides further evidence for the emerging importance of the EGFR signalling pathway in psoriasis and for the first time directly associates this pathway with PsA. These findings may have implication for the treatment of these conditions and may justify repositioning of previously approved EGFR inhibitors, which have been used extensively in cancer chemotherapy (30), for the treatment of psoriasis and PsA. Providing evidence for the roles of the EGFR/ERRFI1 signalling axis in psoriatic disease also furthers our general understanding of these conditions and may open new avenues for study of their pathogenesis.

## Results

### Bioinformatic analysis of 1p36.23 region

The lead SNP for the 1p36.23 risk association – rs11121129 – has previously been shown to be significantly associated with Ps and PsA susceptibility in several GWASs (26-28). To determine a lead causal SNP for the association, a prioritised list of variants was assembled by taking all SNPs in LD with rs11121129 with an R^2^ ≥ 0.8. This comprised a list of 10 variants spanning an interval of ∼13Kb (Table 1). To identify SNPs which may lie in putative regulatory elements, prioritised SNPs were mapped against the genome with key epigenetic markers (ENCODE histone modifications (H3K27Ac, H3K4Me1, H3K4Me3), DNase hypersensitivity sites (DHSs) (31)) using the UCSC genome browser (Figure 1). While rs11121129 did not appear to overlap any key marks of regulatory chromatin, rs11121131 (R^2^=0.9687) overlapped a DHS and peaks of H3K4Me1 and H3K27Ac – the classical signature of an enhancer element (24) – in a number of cell types. Of note, the site is highly enriched for H3K27Ac in the epithelial, endothelial and structural cell types in the ENCODE dataset (Normal Human Epidermal Keratinocytes (NHEK), Human Skeletal Muscle Myoblasts (HSMM) and Human Umbilical Vein Endothelial Cells (HUVEC)) but shows minimal enrichment in bone marrow and lymphoid cell types (K562, GM12878) (Figure 1). This implied that the putative enhancer was active in endothelial and epithelial cells rather than immune cell types. To verify, the variants were also overlayed onto imputed group averages for H3K27Ac and ATAC-seq peaks from the EpiMap primary cell data set (32). Again, the rs11121131 element was enriched for open chromatin and H3K27Ac histone marks in the Epithelial and Endothelial cell sample groups, but not the Blood & T-cell sample group (Additional file 1: Figure S1). ChromHMM chromatin state analysis from the EpiMap dataset also showed that the rs11121131 regulatory element was – on average – represented by highly active chromatin states (Enhancer A1 (EnhA1)) in epithelial and endothelial cell types, while it was most typically represented by inactive or bivalent chromatin states (Bivalent enhancer (EnhBiv) or Weakly repressed polycomb (ReprPCWk)) in blood and immune cell types (Additional file 1: Figure S2). Taken together, these data suggest that rs11121131 may be the causal SNP for the 1p36.23 locus, and that it lies in a putative regulatory element which is active in the epithelium and endothelium rather than immune cell types.

**Figure 1:**
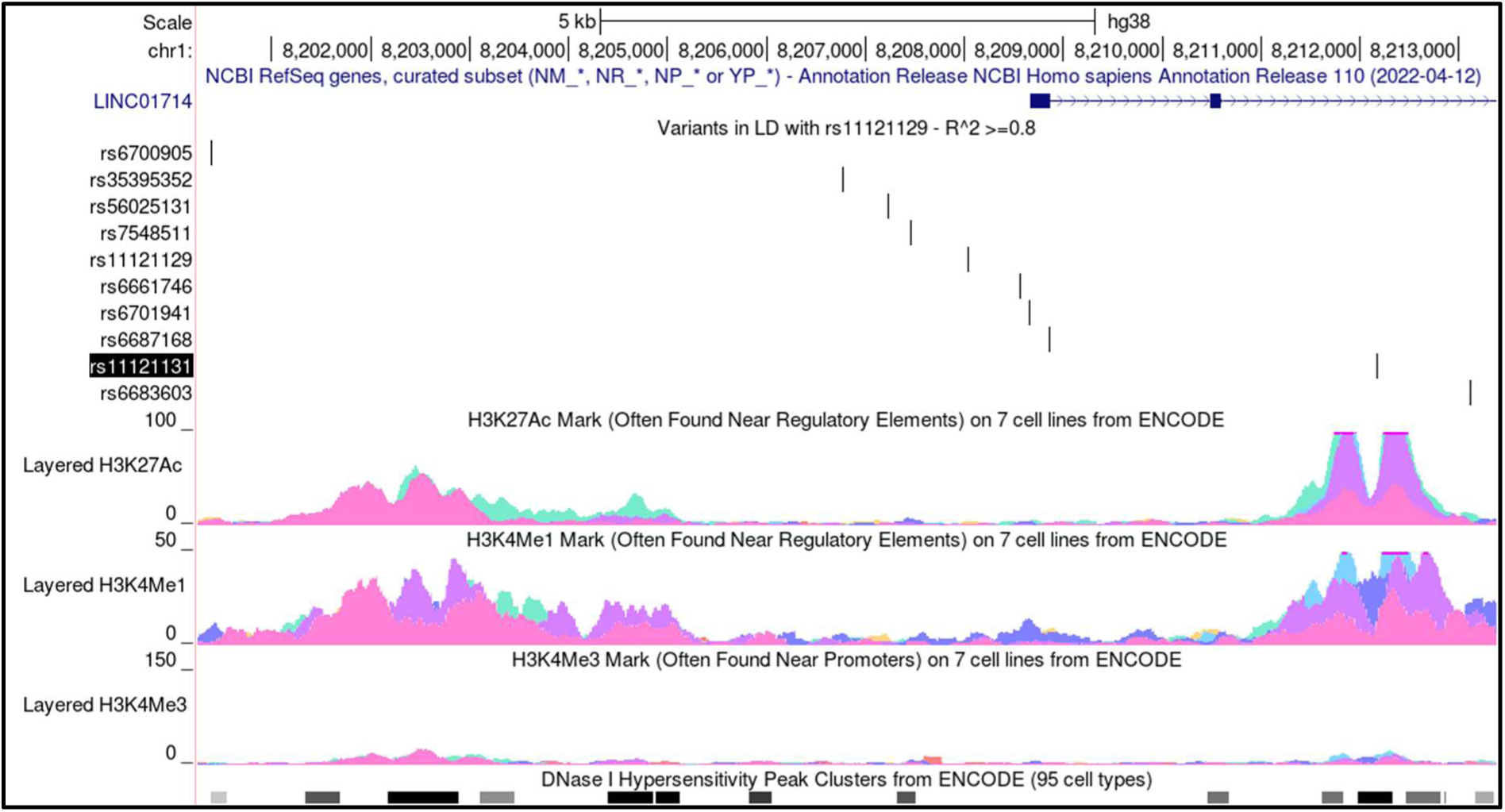
Prioritised SNP variants at 1p36.23. UCSC image showing the 10 variants prioritised for the rs11121129 risk association at 1p36.23, with an R^2^ > 0.8 relative to rs11121129, in their genomic context. The image also shows relevant ENCODE histone and DNA accessibility data.

**Table 1:** Prioritised variants at 1p36.23. Table listing the prioritised SNPs for the 1p36.23 association, made up of all the SNPs in linkage disequilibrium with the lead SNP rs11121129 with an R^2^ score of ≥0.8.

| SNP variant ID | Coordinates (hg38) | Alleles (Major/Minor) | Minor Allele Frequency (MAF) | R<sup>2</sup> relative to rs11121129 |
| --- | --- | --- | --- | --- |
| rs11121129 | chr1:8208034 | (G/A) | 0.2535 | 1 |
| rs6661746 | chr1:8208559 | (G/A) | 0.2535 | 1 |
| rs6701941 | chr1:8208654 | (T/C) | 0.2535 | 1 |
| rs6687168 | chr1:8208852 | (C/T) | 0.2535 | 1 |
| rs7548511 | chr1:8207452 | (A/G) | 0.2555 | 0.9688 |
| rs56025131 | chr1:8207223 | (A/G) | 0.2555 | 0.9688 |
| rs11121131 | chr1:8212170 | (G/A) | 0.2535 | 0.9687 |
| rs35395352 | chr1:8206768 | (-/A) | 0.2565 | 0.9638 |
| rs6700905 | chr1:8200388 | (G/C) | 0.2753 | 0.8744 |
| rs6683603 | chr1:8213116 | (T/G) | 0.2535 | 0.8486 |

Analysis of the rs11121131 element with the JASPAR transcription factor (TF) binding site dataset (Additional file 1: Figure S3) (33), showed the presence of binding sites for transcription factors important in skin and epithelial proliferation and homeostasis, such as the Transcriptional Enhanced Associate Domain (TEAD) family of transcription factors – which are involved in EGFR and Hippo signalling in keratinocytes (34) – and Krüppel-like Factor 4 (KLF4), which our group has previously linked to Ps and PsA in keratinocytes (35). While rs11121131 does not appear to alter any core TF binding motifs, it flanks several predicted TF binding sites. These include E74-Like ETS Transcription Factor 1 (ELF1) – which has suggested involvement in both psoriasis and systemic sclerosis (36, 37) – and ETS Homologous Factor (EHF), which has been implicated as a TF of potential importance in Ps and OA (38, 39). There is evidence to suggest that variants flanking core binding motifs can still significantly affect TF binding, likely by altering the local shape of DNA (40-43), and therefore rs11121131 may still have some functional impact on the binding of these TFs.

To identify putative target genes for this locus, we analysed the locus in previously published promoter capture Hi-C and H3K27Ac HiChIP datasets from our group (35, 44). These experiments were carried out in both the HaCat keratinocyte cell line and the MyLa CD8^+^ T cell line. Both Hi-C and H3K27Ac HiChIP showed that – in HaCat cells – looping could be observed between the rs11121131 regulatory locus and a gene locus ∼200Kb away containing 3 genes: Tumour Necrosis Factor Super Family 9 (*TNFRSF9*), Parkinsonism Associated Deglycase (*PARK7*) and ERBB receptor feedback inhibitor 1 (*ERRFI1*) (Figure 2). All 3 of these genes are of potential functional interest in psoriatic disease. *TNFRSF9* is a key co-stimulatory receptor in activated T-cells (45), *ERRFI1* is a negative regulator of EGFR signalling, which is a major driver of proliferation and differentiation in keratinocytes and other related cell types (46, 47), and *PARK7* has potentially important roles in cell cycle control and oxidative stress response (48-50).

**Figure 2:**
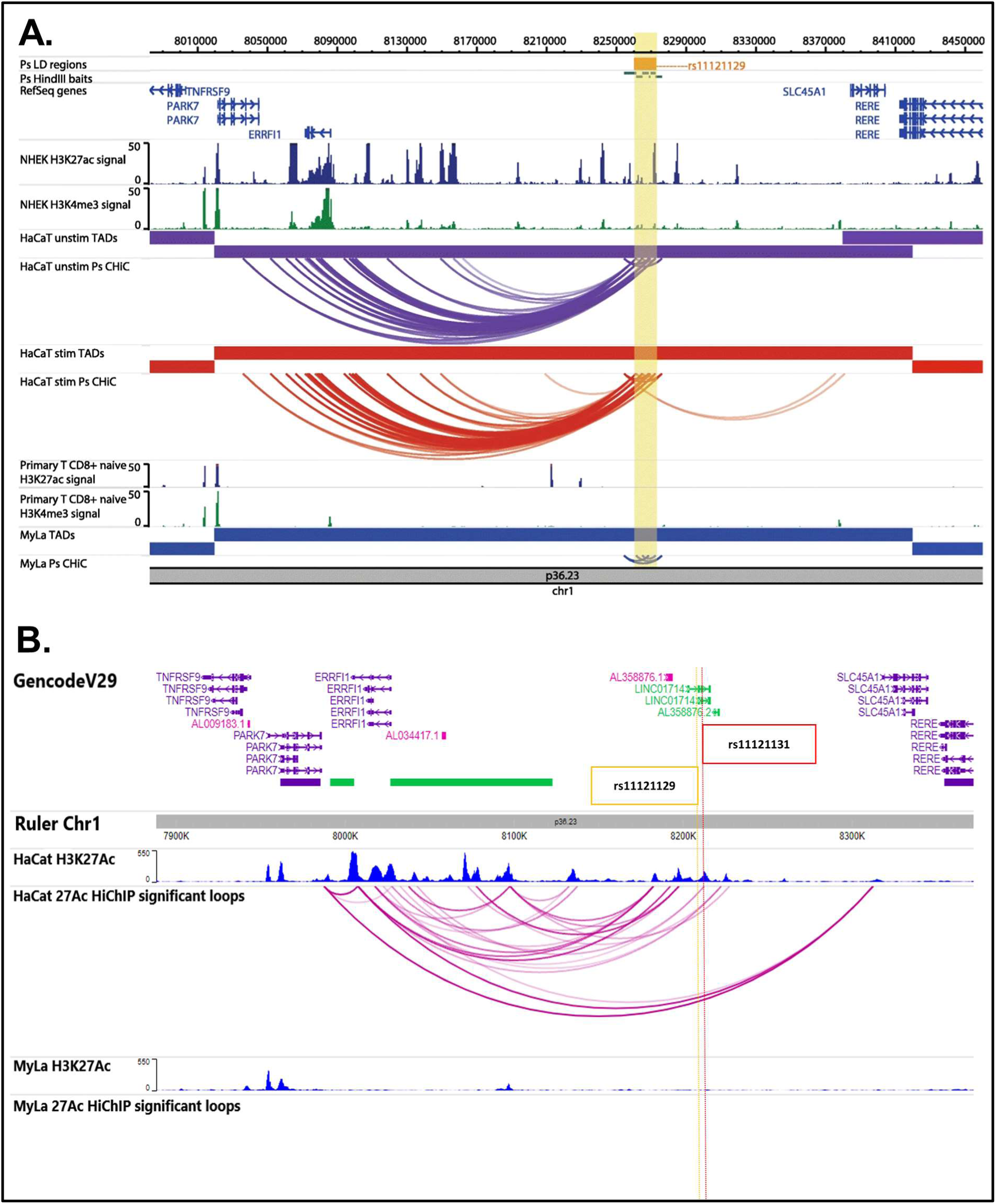
Capture Hi-C and HiChIP show keratinocyte-specific interactions at PsA/psoriasis risk region 1p36.23. **A.**Figure taken from Ray-Jones et al., 2020 (35) (original publisher: BMC Biology) showing interactions between the 1p36.23 risk locus and the TNFRSF9/PARK7/ERRFI1 gene locus in the HaCat keratinocyte cell line. This interaction is not present in the MyLa CD8+ T-cell line, which suggests that this interaction is important in keratinocytes and related cell types, rather than immune lineage cells. **B.** WashU browser image showing H3K27Ac ChIP peaks the HiChIP interaction loops called by FitHiChIP from HaCat and MyLa cells in their genomic context. The location of the rs11121131 putative enhancer element has been marked. As can be seen, the same interactions between the rs11121129/rs11121131 risk locus and the TNFRSF9/PARK7/ERRFI1 gene locus observed in the Capture Hi-C data are present again for HaCat cells. Again, no interactions were observed in MyLa cells.

Interestingly, this chromatin looping between the 1p36.23 risk region and the *TNFRSF9/PARK7/ERRFI1* locus was not observed in MyLa cells, further supporting the non-lymphoid nature of this risk locus. In the H3K27Ac HiChIP experiment in particular – which captures loops between active regions of chromatin (51) – looping was observed between H3K27Ac peaks at the rs11121131 element and the promoter of *ERRFI1* (Figure 2B). No enrichment of H3K27Ac or chromatin looping was observed at the 1p36.23 region in MyLa cells. It is also of note that rs11121131, along with the other SNPs in tight LD, is an expression Quantitative Trait Locus (eQTL) for *ERRFI1* in cultured human fibroblasts (GTEx V8, P = 4.4×10^-13^), which indicates that being homozygous for the minor allele of rs11121131 leads to decreased levels of *ERRFI1*. This eQTL prioritises *ERRFI1* as a candidate causal gene for the 1p36.23 risk association, but requires validation as to causal SNP, enhancer, cell type and mechanism. We therefore applied a range of CRISPR-Cas9 based techniques to functionally dissect the locus to determine both the true causal regulatory element and causal gene for this risk locus.

### CRISPR-Cas9 mediated deletion of putative regulatory regions within 1p36.23 in HaCat keratinocytes

Using classic cut-and-repair CRISPR-Cas9 gene editing we deleted the rs11121131 element in HaCat cells and measured the impact on surrounding gene activity.

Based on the ENCODE H3K27Ac signal from NHEK cells we decided to delete a 2.1kb region – 1kb either side of rs11121131 – to remove the whole putative regulatory element (Figure 3A). We designed single guide RNAs (sgRNAs) on either side of the region to be deleted, and both sgRNAs were delivered as *Streptococcus pyogenes* Cas9 (SpCas9) ribonucleoprotein (RNP) complexes together with a plasmid homology donor template to replace the intervening region with a Cre-excisable puromycin resistance cassette via homology-directed repair (HDR) (Additional file 1: Figure S4A). After selection of edited cells via puromycin, transduction of cells with an AAV vector expressing Cre recombinase and GFP allowed removal of the puromycin cassette and single-cell sorting of transduced cells. This results in the deleted region being replaced by 134bp of sequence containing a single loxP site. The other allele in the cells may be repaired by a second HDR event or simply by non-homologous end joining (NHEJ) driven deletion between the two sgRNAs.

**Figure 3:**
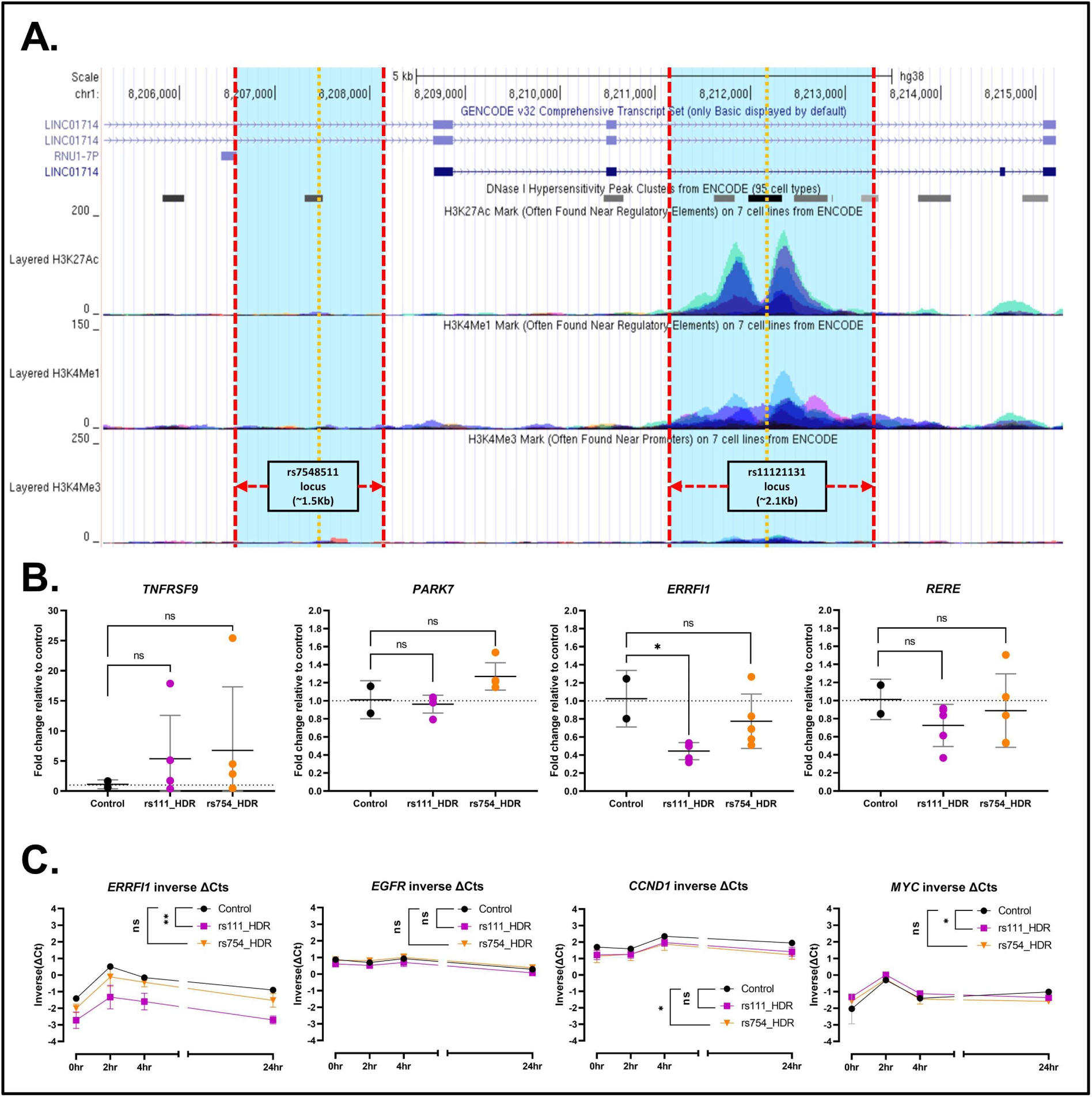
Deletion of putative functional regions of 1p36.23 in HaCat cells. **A.**Locations of the deleted regions overlayed on UCSC (GRCh38/hg38), showing ENCODE histone marks and DNA accessibility data. The location of the SNP of interest for each site is indicated by the vertical yellow lines. **B.** RT-qPCR analysis of steady-state deletion cells (n=5) compared to normal polyclonal HaCat controls (n=2) for the four potential target genes of the 1p36.23 risk locus. Data analysed via one-way ANOVA with a post-hoc Tukey’s test to determine significance relative to control cells. **C.** RT-qPCR analysis of EGF stimulation time course for 4 EGF-responive genes. Deletion clones (n=3) were compared to normal polyclonal HaCat cells (n=3) at 0, 2, 4 and 24hrs post stimulation. Data analysed using a linear mixed effects model (nlme package in R), with significance shown for the fixed effect of deletion class (control, rs111_HDR or rs754_HDR) on inverse(ΔCt) values (calculated by −1*(ΔCt)) across the whole timecourse. For qPCR, 20ng RNA per assay, all assays carried out in triplicate. Significance summaries – ns: P≥0.05; *: P<0.05; **: P<0.01; ***: P<0.001; ****: P<0.0001.

For comparison, a region surrounding another SNP – rs7548511 – was selected for deletion (Figure 3A), as this SNP does not lie in a putative regulatory element.

6 single cell clones were selected for the rs11121131 deletion (rs111_HDR) and 5 single cell clones for the rs7548511 deletion (rs754_HDR). PCR to detect amplicons in the deleted region showed that 5/6 rs111_HDR clones had biallelic deletions, as did all 5 rs754_HDR clones (Additional file 1: Figure S4B). In each case, the samples were also assayed for the non-deleted region to help control for large deletions which may have removed both SNP regions. Removal of the Puromycin selectin cassette was confirmed by PCR. In all cases where biallelic deletion was observed, no amplicon of the size corresponding to an intact puromycin resistance cassette was observed (Additional file 1: Figure S4C).

The 5 homozygous deletion clones for each region were taken forward for Reverse-transcription quantitative PCR (RT-qPCR) analysis. Cells were assayed for expression of *TNFRSF9*, *PARK7*, *ERRFI1* and *RERE* – the closest gene on the 3’ side of the 1p36.23 region. Solute Carrier Family 45 Member 1 (*SLC45A1*), another nearby gene, was discounted as it was undetectable by RT-qPCR in HaCat cells. Compared to normal HaCat cells, cells with the rs11121131 region deleted exhibited a significant ∼56% mean decrease in levels of *ERRFI1* mRNA across the 5 clones, which was not observed with the rs7548511 deletion (Figure 3B). No significant effects were observed on the other 3 genes. It should be noted that *TNFRSF9* exhibited large, inconsistent changes in expression between clones. This is likely due to a low basal mRNA level of this gene, leading to small clonal differences in expression appearing more pronounced. This therefore is strong evidence that the enhancer containing the risk variant rs11121131 has a significant impact on the expression of *ERRFI1*, and more so than the rs7548511 deletion.

As *ERRFI1* is a regulator of the epidermal growth factor (EGF) receptor, we decided to see how deletions affected expression of EGF-responsive gene across a time course when cells were stimulated with EGF. 3 rs111_HDR clones and 3 rs754_HDR clones were compared to normal polyclonal HaCat cells at 0, 2, 4 and 24hrs post-stimulation with 10ng/mL recombinant EGF. Clones where two different alleles could be seen on the agarose gel analysis were selected, as it was thought these were less likely to harbour unintended larger deletions. Cells were assayed by RT-qPCR for expression of *ERRFI1* and *EGFR*, along with two EGF-responsive cell cycle regulators: *MYC* and *CCDN1*. Data were analysed using a linear mixed effects model (nlme package in R), with fixed effects of deletion class (WT, rs111_HDR or rs754_HDR), time, and the interaction of deletion class and time. Sample ID was included as a random effect to minimise the impact of clonal differences not arising from the deletions. The rs11121131 deletion, but not the rs7548511 deletion, was shown to significantly lower expression of *ERRFI1* and significantly alter expression of *MYC* across the stimulation time course (Figure 3C; Additional file 1: Figure S5). Although only the rs11121131-deleted clones impacted the expression of *ERRFI1* across the stimulation time course, deletion of rs7548511 did significantly alter expression of *CCND1* in the time course, which may point to some regulatory involvement. When the interaction effect of deletion class and time was analysed, it was discovered that both deletions significantly alter the way *MYC* expression changed between the 0hr and 24hr mark, which may again point to deletion of the rs7548511 region having some regulatory consequences, perhaps as a co-operative element in the control of the rs11121131 element or by altering local chromatin structure. For the time course samples, we also calculated the fold change in mRNA levels relative to controls at the 0hr timepoint for all genes. It was observed that both the rs11121131 deletion and the rs7548511 deletion significantly reduced levels of *ERRFI1*, but other genes were not significantly altered at 0hr (Additional file 1: Figure S6).

Taken together, these data suggest that the rs11121131 regulatory element contributes to the regulation of *ERRFI1* expression. Some regulatory changes were also observed upon deletion of the rs7548511 element in the time course, and this suggests it may also have regulatory roles affecting *ERRFI1* in certain circumstances, for instance here only detected 24hrs post-stimulation. Regardless, these observed effects were neither as pronounced nor as consistent as those observed with the rs11121131 element, positioning this putative enhancer as the likely driver of *ERRFI1* regulation in the region.

### Perturbations of the 1p36.23 locus via CRIPSR activation and interference

As a complementary assay, we decided to perturb rs11121131 element and the rs7548511 region via CRISPR activation (CRIPSRa) and CRISPR interference (CRISPRi) in HaCat cells. CRISPRa and CRISPRi tools can be used to activate or repress regulatory elements have been shown to also affect their regulated genes, providing a functional link between the enhancer and its target (24).

Firstly, the two loci were targeted using a previously-validated HaCat line stably expressing dCas9-p300 and a puromycin resistance gene (35). 3 sgRNAs were designed per locus, either spanning the putative regulatory region in the case of the rs11121131 locus or similarly spaced around the rs7548511 SNP (Additional file 1: Figure S7). In both cases, all 3 sgRNAs were designed to target the region deleted in previous experiments. sgRNAs were cloned into a lentiviral delivery vector co-expressing a Neomycin resistance gene (pLKO5.sgRNA.NeoR), to allow antibiotic co-selection of cells stably expressing the activator and the sgRNA. On-target sgRNAs were compared to two non-targeting sgRNA controls - the unmodified pLKO5.sgRNA.NeoR vector, in which there is a non-targeting spacer sequence to act as a placeholder for cloning (referred to as pLKO5.NeoR (empty) from here forward) and scrambled control sgRNA from the literature (“Scramble2”) (52).

We observed that targeting dCas9-p300 to the rs11121131 element led to significant upregulation of *ERRFI1* (Figure 4), re-affirming the functional relationship between this putative enhancer and the gene. Similar upregulation of *ERRFI1* was also observed upon targeting the rs7548511 site (Figure 4), adding further support to the idea this may be a co-regulatory element at the 1p36.23 locus affecting *ERRFI1*.

**Figure 4:**
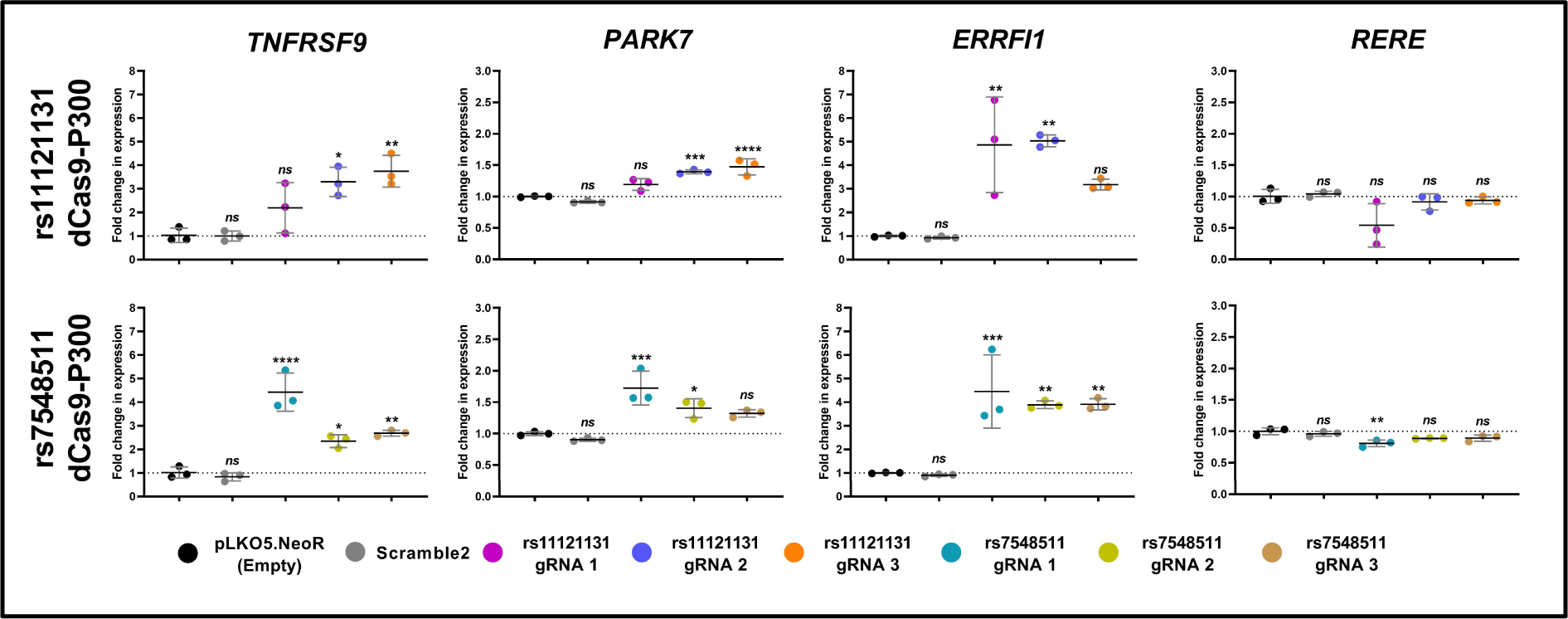
Activation of rs11121131 and rs7548511 loci with dCas9-p300. RT-qPCR results from CRISPRa experiments in HaCat cells expressing dCas9-p300 using gRNAs targeted to rs11121131 or rs7548511, alongside pLKO5.sgRNA.NeoR empty vector (pLKO5.NeoR (empty)) and Scramble2 controls. n=3 biological replicates per condition. For RT-qPCR, 20ng RNA was used per assay, and all assays were carried out in triplicate. Fold change was calculated relative to the mean of the pLKO5.NeoR (empty) controls via the 2^-ΔΔCt^ method. Data analysed via one-way ANOVA with a post-hoc Tukey’s test to determine significance relative to pLKO5.NeoR (empty) control (ns: P≥0.05; *: P<0.05; **: P<0.01; ***: P<0.001; ****: P<0.0001).

Interestingly, we also observed milder upregulation of both other genes at this site – *TNFRSF9* and *PARK7* – in contrast to the enhancer deletion experiment which showed no effect on the activity of these genes. We hypothesised this may be a result of the use of dCas9-p300, which is a direct histone acetylase. As previous Capture Hi-C data shows a clear interaction between the 1p36.23 locus and the *TNFRSF9*/*PARK7*/*ERRFI1* locus, dCas9-p300 is possibly hyper-acetylating this region, thus driving expression of the other genes. This would also explain why we observed no significant effect on *RERE*, as no interactions between this gene and 1p36.23 were observed in the conformation capture assays. Indeed, others have seen similar non-specific regulatory effects when utilising dCas9-p300 (53). As an alternative to dCas9-p300 we generated a new HaCat activator cell line stably expressing an alternative CRISPRa system: dCas9-VPR, a potent transcriptional activator complex which functions as a scaffold to recruit endogenous regulatory proteins (54), rather than acting as an direct epigenetic modifier like dCas9-p300. After validation of the activation potential of this cell line using a positive control sgRNA (Additional file 1: Figure S8), HaCat-dCas9-VPR cells were transduced with the same sgRNAs used in the dCas9-p300 experiment.

With the VPR system, no significant effects were observed on the expression of *TNFRSF9*, *PARK7* or *RERE* with the targeting sgRNAs, but there was a significant 35.8% to 48.4% decrease in *ERRFI1* transcript levels across all 3 sgRNAs at the rs11121131 locus, and a significant 46.7% to 53.6% decrease was observed across the 3 sgRNAs targeting the rs7548511 locus (Figure 5A). Small but significant differences between the two control sgRNAs for *ERRFI1* and *PARK7* were also observed. For *ERRFI1*, the observed change was in the opposite direction-of-effect to the observed effects of the on-target sgRNAs, and the downregulation of *ERRFI1* remains highly significant compared to either control. For *PARK7*, the observed increase was small (1.13-fold), and unlikely to be of biological significance within the context of this assay.

**Figure 5:**
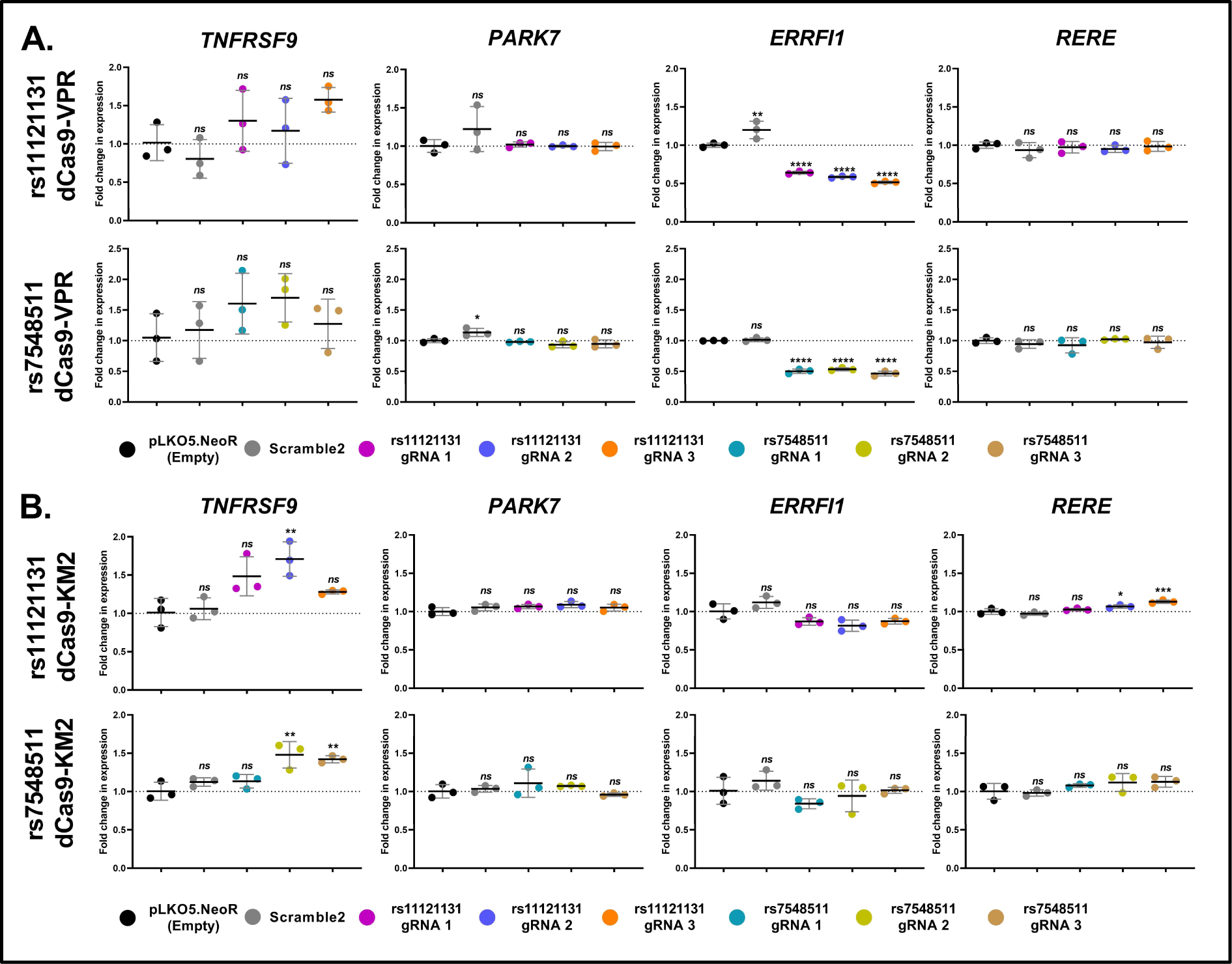
CRISPRa and CRISPRi at the 1p36.23 locus with dCas9-VPR and dCas9-KM2. RT- qPCR results from experiments in HaCat cells targeting the rs11121131 and rs7548511 loci with **A.** dCas9-VPR or **B.** dCas9-KRAB-MeCP2 (dCas9-KM2), alongside pLKO5.sgRNA.NeoR empty vector (pLKO5.NeoR (empty)) and Scramble2 controls. n=3 biological replicates per condition. For RT-qPCR, 20ng RNA was used per assay, and all assays were carried out in triplicate. Fold change was calculated relative to the mean of the pLKO5.NeoR (empty) controls via the 2^-ΔΔCt^ method. Data analysed via one-way ANOVA with a post-hoc Tukey’s test to determine significance relative to pLKO5.NeoR (empty) control (ns: P≥0.05; *: P<0.05; **: P<0.01; ***: P<0.001; ****: P<0.0001).

The enhancer deletion experiments resulted in a reduction in *ERRFI1* expression, CRISPRa using the p300 epigenetic modifier led to upregulation of *ERRFI1*, whereas the CRISPRa experiment using dCas9-VPR paradoxically results in repression of this gene. Taken together, these results confirm the regulatory target of the 1p36.23 risk locus as *ERRFI1*.

As a complementary assay, a HaCat line expressing the CRISPR repressor dCas9-KRAB-MeCP2 (dCas9-KM2) (55) was also developed and validated with a positive control sgRNA (Additional file 1: Figure S8). Effects observed with repression of the two elements were generally minor or insignificant, with no significant effects observed upon *PARK7* or *ERRFI1* (Figure 5B). However, some significant increases in *TNFRSF9* expression were observed. While it should be observed that *TNFRSF9* is only detectable at low levels in HaCat cells, making accurate observations by RT-qPCR difficult, these may indicate some regulatory effect on *TNFRSF9* at this locus. There were also some small but significant increases in the expression of *RERE* observed when targeting rs11121131, but in the context of this assay changes of this magnitude are unlikely to be of biological significance.

Collectively, these data indicate *ERRFI1* as a regulatory target of the 1p26.23 risk locus, but also indicate a complex mechanism of regulation.

### Histone ChIP-qPCR for markers of activation and repression in HaCat-dCas9-VPR cells activating 1p36.23

The stark differences between CRISPRa experiments using different effectors (p300 leading to activation, and VPR to repression) indicated a potential role of histone modification in the function of the enhancer. We performed ChIP-qPCR analysis of regions at both the *ERRFI1* promoter – both at the proximal promoter sequences of the main isoform (hereafter the main promoter) and also the sequences upstream of an annotated short isoform of the gene, which initiates partway through exon 2 – and the rs11121131 and rs7548511 SNP loci. ChIP was carried out for both H3K27Ac – a marker of active chromatin – and H3K27Me3, a reciprocal marker of transiently repressed chromatin (Figure 6). For both rs11121131 and rs7548511, the sgRNA which had the most pronounced repressive effect was selected for ChIP-qPCR study, which was sgRNA 3 in each case. Negative control immunoprecipitations were also carried out using an IgG control antibody (Additional file 1: Figure S9).

**Figure 6:**
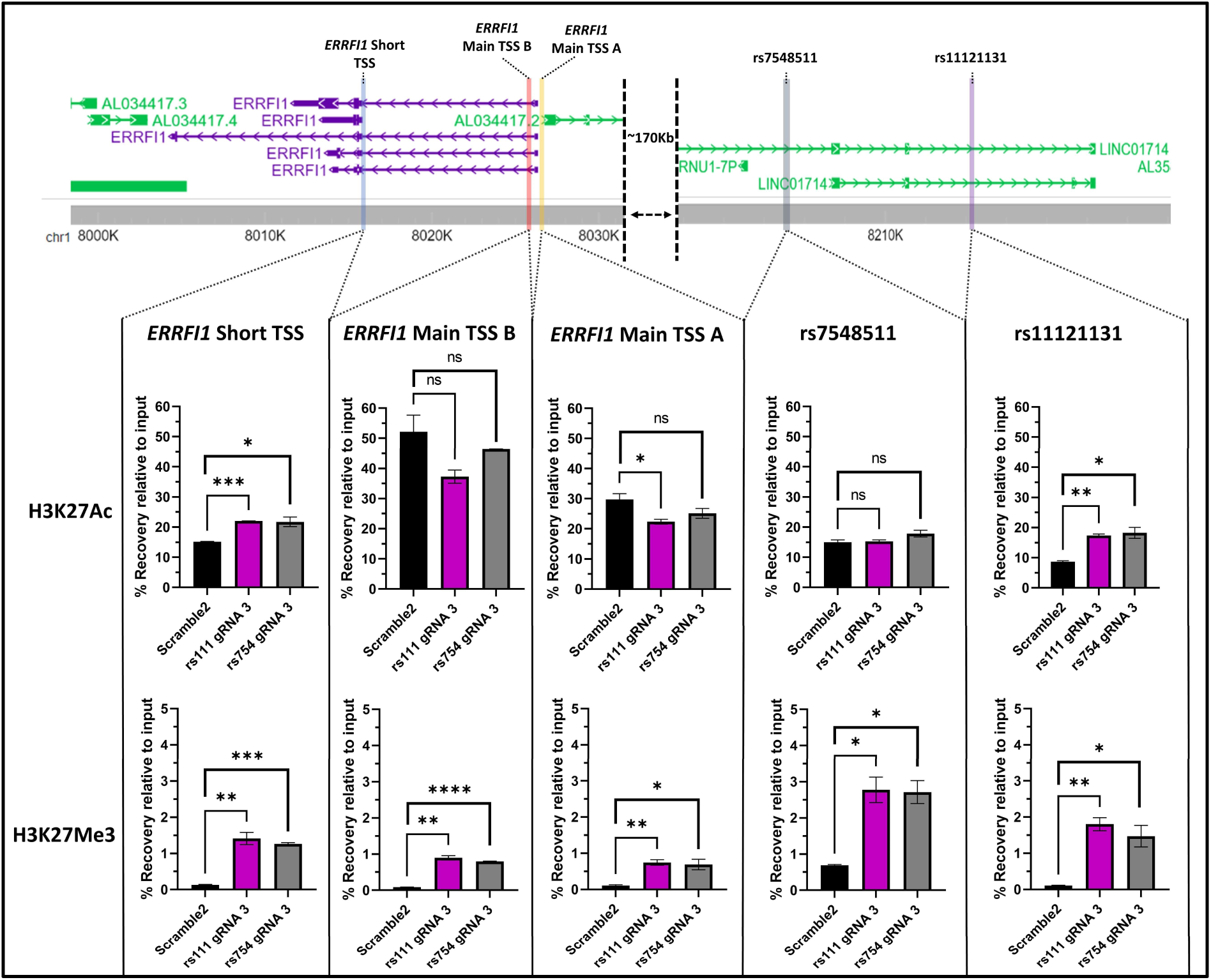
Histone ChIP-qPCR in HaCat-dCas9-VPR cells targeting rs11121131 or rs7548511. Diagram detailing the results of the H3K27Ac and H3K27Me3 histone ChIP-qPCR experiments in the HaCat-VPR cell lines, expressing either a scrambled control gRNA (Scramble2), a gRNA targeting the rs11121131 SNP locus (rs111 gRNA 3) or a gRNA targeting the rs7548511 gRNA locus (rs754 gRNA 3). The top panel shows the relative location of each qPCR primer pair on the genome, with the bottom panels showing the percentage recovery of the locus for each histone mark in each cell line. n=2 biological replicates for each condition. qPCR was carried out using 1μL of sample per 10μL reaction using the BioRad SsoAdvanced universal SYBR green Supermix, with 3 technical replicates per reaction. Statistical testing was carried out by individual Student’s t-tests (ns: P≥0.05; *: P<0.05; **: P<0.01; ***: P<0.001; ****: P<0.0001). For IgG negative controls, see Additional File 1: Figure S9. For qPCR primers used, see Additional file 2: Table S3.

dCas9-VPR targeted to either SNP only significantly increased H3K27Ac expression at the rs11121131 SNP locus. This suggests that the regulatory effect observed from targeting rs7548511 was likely being driven through co-operative activation of the rs11121131 enhancer, either through a true regulatory mechanism or an artificial effect of the activator. Regardless, these data again point to the rs11121131 locus being the key regulatory element at 1p36.23. At the *ERRFI1* locus, a small significant decrease was observed at one of the main *ERRFI1* promoter sites when rs11121131 was activated, but no other significant changes in H3K27Ac were observed at the *ERRFI1* main promoter. Interestingly, a significant increase in H3K27Ac was observed at the short isoform promoter of *ERRFI1*. This isoform is not well described; however it should be noted that the TaqMan qPCR assay used for *ERRFI1* detection throughout this work does not target this isoform.

When H3K27Me3 signal was assayed, a large and significant deposition of H3K27Me3 marks could be observed across all target sites when dCas9-VPR was targeted to either the rs7548511 or the rs11121131 loci, substantially altering the ratio of H3K27Ac:H3K27Me3 (Table 2). This is consistent with the repressive effect observed in the dCas9-VPR experiment. These results, taken together with the CRISPR activation and interference experiments, point to a more complex mechanism of regulation between the 1p36.23 risk locus and *ERRFI1*, and one which warrants further study.

**Table 2:** H3K27Ac:H3K27Me3 ratios from ChIP-qPCR experiments. Table showing the ratio of H3K27Ac recovery to H3K27Me3 recovery for each cell line at each of the loci analysed by ChIP-qPCR. Ratios were calculated by dividing the average H3K27Ac percentage recovery by the average H3K27Me3 percentage recovery for each target site.

| ChIP target locus | Scramble2 control<br>H3K27Ac:H3K27Me3<br>ratio | rs111 gRNA 3<br>H3K27Ac:H3K27Me3<br>ratio | rs754 gRNA 3<br>H3K27Ac:H3K27Me3<br>ratio |
| --- | --- | --- | --- |
| rs11121131 SNP locus | 81.57 | 9.65 | 12.39 |
| rs7548511 SNP locus | 21.59 | 5.51 | 6.57 |
| *ERRFI1* Main TSS A | 275.90 | 29.95 | 36.48 |
| *ERRFI1* Main TSS B | 618.23 | 41.57 | 57.79 |
| *ERRFI1* Short TSS | 118.18 | 15.60 | 17.29 |

### Assessing the role of *ERRFI1* in skin inflammation *in vivo*

We sought to investigate whether *ERRFI1* played a role in psoriatic skin inflammation *in vivo* by utilising the imiquimod-induced psoriasis-like mouse model (56). Psoriasis-like plaque formation was induced in C57BL/6 mice by topical application of 10mg Aldara cream (5% imiquimod W/W) to the ear. Onset of inflammation and development of psoriasis-like plaques were confirmed via scoring of skin thickening, scaling and redness, as well as flow cytometry analysis of immune cell infiltration and IL-17 production (Additional file 1: Figure S10). Samples of inflamed skin were then taken from these mice for histological analysis to detect MIG-6 – the mouse homolog of ERRFI1 – alongside non-inflamed controls.

Immunofluorescent staining of ear skin sections from inflamed and non-treated animals revealed that inflamed tissue exhibited increased staining for MIG-6 in the epidermal layer when compared to non-treated controls (Figure 7; Additional File 1: Figures S11-S15). Co-staining for cytokeratin was used to confirm that this increased staining was observed in epidermal keratinocytes, and samples were also stained with isotype controls to confirm the specificity of the MIG-6 staining.

**Figure 7:**
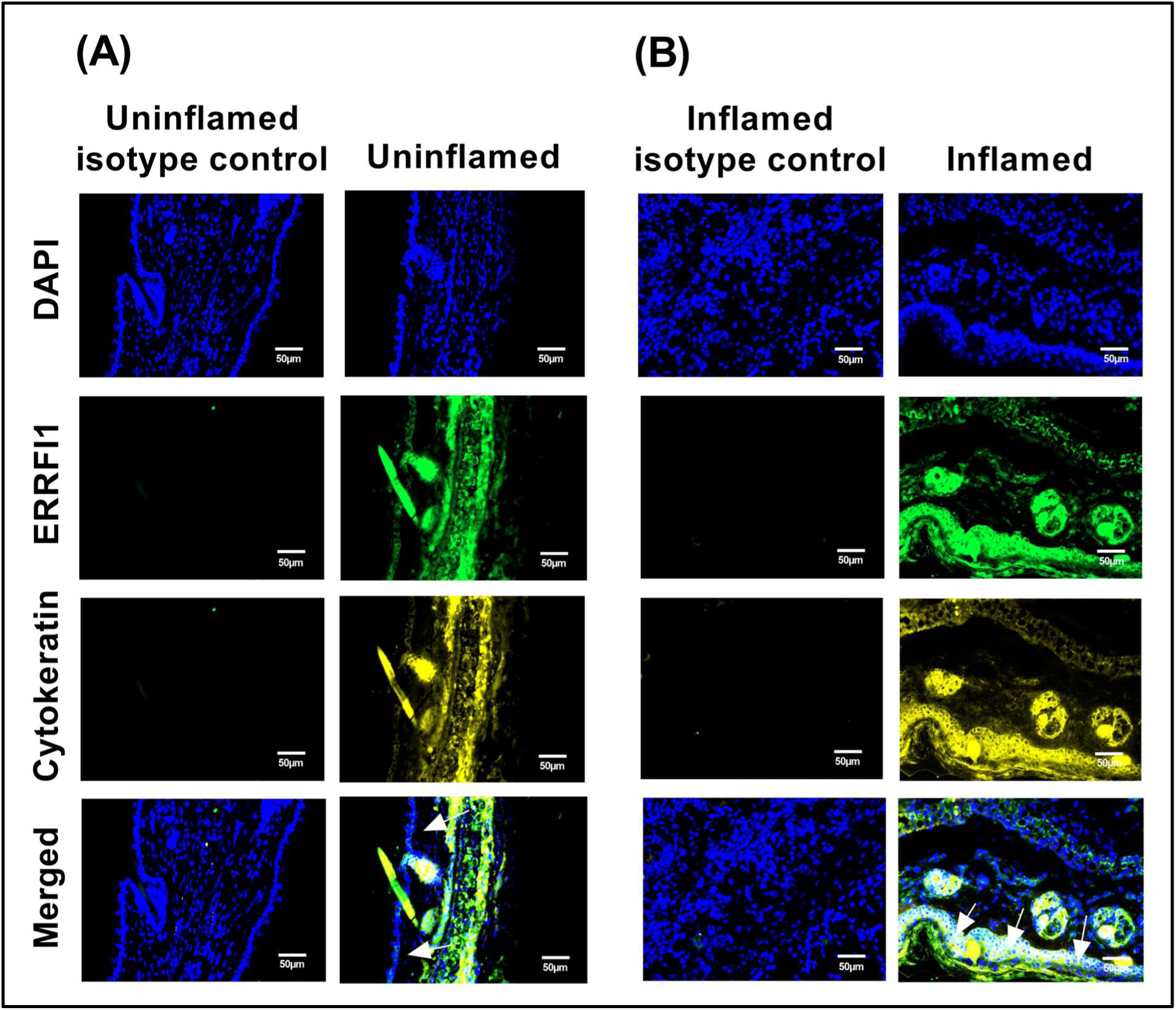
ERRFI1 is expressed in inflamed epidermal keratinocytes in the murine Ps model. Immunofluorescent imaging of sections taken from the ear skin of imiquimod-induced psoriasis-like mouse models (**B**) vs uninflamed controls (**A**). C57BL/6 mice were treated topically with Imiquimod-containing Aldara cream daily for 3 days to induce psoriasis-like skin inflammation. C57BL/6 mice ear tissues were harvested, cryo-sectioned and stained for ERRFI1 expression, alongside cytokeratin 19 to help identify the epidermal keratinocyte layer and a DAPI nuclear co-stain. Controls were taken from untreated C57BL/6 mice. The epidermal layer in the inflamed skin sample (**B**, white pointers) appears to show a considerable increase in ERRFI1 staining compared to the control samples (**A**, white pointers), accompanying the epidermal hyperplasia typical of psoriasis-like inflammation. Appropriate isotype controls were used to confirm the specificity of the observed staining. Scale bars show 50µm. Data is representative of n=6 treated mice vs 6 untreated. Immunofluorescence staining from these additional mice can be found in Additional File 1: Figures S11-S15.

The increased MIG-6 staining in epidermal keratinocytes in response to psoriasis-like inflammation suggest an upregulation of MIG-6 during the progression of psoriatic plaque formation. These data provide evidence for the involvement of ERRFI1 in psoriatic inflammation. When considered in light of our GWAS and *in vitro* investigations – which suggest a genetic variant in a psoriasis-associated risk region modulates the expression of *ERRFI1* – these data strongly suggest a novel role for *ERRFI1* in the risk of psoriatic disease.

## Discussion

The data presented here show that rs11121131 – a putative causal variant for the 1p36.23 Ps and PsA risk association – lies in an enhancer element which controls the expression of *ERRFI1*. While this regulatory region may have additional functions on other genes, perhaps indicating a role as a multi-gene enhancer, there is a clear and consistent effect on *ERRFI1*, strongly suggesting it is the main regulatory target of this risk locus.

Multiple strands of evidence suggest *ERRFI1* may be important in the development of Ps and PsA. *ERRFI1* knock-out mouse models exhibit skin phenotypes indicative of psoriasis, such as impaired differentiation and hyperplasia of keratinocytes (57, 58). Furthermore, joint-restricted knockout and overexpression of *ERRFI1* in mouse models led to osteoarthritis-like phenotypes (59-61). It has also recently been shown that EGFR signalling – of which ERRFI1 is a key regulator – can enhance cytokine-driven IL-23 production in keratinocytes (62), a key inflammatory modality in Ps. Thus, ERRFI1 and the EGFR signalling pathway are key to joint and skin homeostasis, and that disruption can produce phenotypes indicative of disease.

To provide further evidence for the role of ERRFI1 in psoriatic skin inflammation, we examined protein levels of the mouse ERRFI1 homolog MIG-6 in inflamed skin samples from Imiquimod-induced psoriasis mouse models. We observed increased staining for MIG-6 in the epidermis of inflamed animals, suggesting an upregulation of this protein during the progression of psoriasis. Furthermore, the role of MIG-6/ERRFI1 as a negative feedback regulator of EGFR – and the fact that it is an early response gene for EGF signalling - further suggests increased levels of EGFR activation and signalling in the inflamed skin, driving the increased levels of ERRFI1. When we consider our experiments deleting the rs11121131 enhancer – which led to reduced *ERRFI1* expression – along with the direction-of-effect of the GTEx eQTL observed for rs11121131 upon *ERRFI1*, it is likely that the risk variant at 1p36.23 reduces levels of *ERRFI1*, thereby reducing the ability of ERRFI1 to inhibit EGFR signalling and leading to greater degrees of inflammation. However, in this study we were not able to directly assay the effect of different alleles of rs11121131 upon *ERRFI1* expression, and therefore it is possible that rather than disrupting the enhancer in which it sits, rs11121131 may have a gain-of-function effect. However, given the known regulatory role of ERRFI1 upon EGFR, and the emerging role of EGFR signalling in psoriasis, the mechanism proposed above is still the most likely driver of psoriasis risk.

*ERRFI1* and the EGFR pathway have not previously been linked to the pathogenesis of PsA. For psoriasis, a study by Kubota and Suyama utilised public bioinformatic datasets to propose a link between *ERRFI1* and another SNP at 1p36.23 - rs72635708 – suggesting the SNP modulated AP-1 binding (63). Their SNP is in tight LD with a SNP identified in a meta-analysis between European and Chinese population carried out by Yin et al. (64), and is only in moderate LD with rs11121131 (R^2^ = 0.6331 in European populations, as calculated by LDProxy). It may be that this is another regulatory mechanism at the 1p36.23 locus, as it is likely that there are other regulatory elements which control *ERRFI1* expression, perhaps in response to different stimuli. Regardless, while their study relies on bioinformatic approaches alone, our study provides direct functional evidence of the link between the rs11121131 enhancer element and *ERRFI1* expression. *ERRFI1* has also been linked to Non-Alcoholic Fatty Liver Disease (NAFLD) (65) and control of glucose metabolism (66). These are both hallmarks of metabolic syndrome – the major co-morbidity of psoriatic disease – and suggest that the ERRFI1/EGFR regulatory axis may contribute to the link between these conditions.

An interesting observation of our study was that the region surrounding the SNP rs7548511 – which was initially chosen for comparison as it did not overlap any predicted regulatory elements – appeared to exhibit some regulatory effects upon *ERRFI1*, although these effects were milder and less consistent compared to perturbations of the rs11121131 locus. This result was unexpected, as the region immediately surrounding rs7548511 does not overlap any of the histone marks or ATAC-seq peaks normally associated with regulatory DNA in any of the sample groups from the EpiMap dataset. In the case of the CRISPRa data, ChIP-qPCR revealed that targeting either rs11121131 or rs7548511 led to H3K27 acetylation at rs11121131. This suggests that the regulatory effect of targeting rs7548511 with dCas9-VPR upon *ERRFI1* is likely facilitated through co-operative activation of the rs11121131 enhancer. In the case of the CRISPR deletion experiments, it is possible that deletion of the rs7548511 element also affects the regulatory properties of the rs11121131 enhancer, via altering the local chromatin environment and DNA structure (67). It is also possible that the region surrounding rs7548511 represents a secondary member of an enhancer chain, its activity augmenting that of the main rs11121131 enhancer (68). From these data, therefore, we cannot completely rule out the region surrounding rs7548511 as an independent or co-operative regulatory element. While the vast majority of cis-regulatory elements are marked by open chromatin, H3K27Ac histone marks or both, it has been shown that there can be rare exceptions to this rule (69). However, the rs11121131 enhancer both has the typical epigenetic profile of an active enhancer in keratinocytes, and exhibits a clear and reproducible effect on *ERRFI1* mRNA levels when perturbed. Therefore, it is most likely that the rs11121131 enhancer is the main driver of the Ps and PsA risk association at 1p36.23.

While we have provided strong evidence for the putative causal variant and target gene for the 1p36.23 risk association, the discrepancies in the direction-of-effect of regulation observed between the deletion experiment and the CRISPRa experiments point to a more complex regulatory mechanism at this locus which warrants further study. It is likely that the repressive effect and wide-spread deposition of H3K27Me3 observed in the dCas9-VPR experiments is due to some downstream negative feedback mechanism; a consequence of the steady-state activation of the locus. This mechanism may be better perturbed in future work using an inducible CRISPRa system, to allow temporal control of activation of the locus. We also hypothesise that the repression observed may be a consequence of activation of the long non-coding RNA at the 1p36.23 locus – *LINC01714* – as examples exist in the literature of non-coding RNAs exhibiting *cis*-repressive effects via recruitment of PRC2 and deposition of H3K27Me3 (70, 71). Important next steps would be to model the different variants of rs11121131 – either *in vitro* in cultured cells or *in vivo* in engineered mouse models – to determine the true mechanistic impact of this variant.

While this study was not able to elucidate the full regulatory mechanism of the 1p36.23 locus, or perturb the rs11121131 variant directly, the clear and reproducible effects of perturbing the rs11121131 enhancer on *ERRFI1* – combined with the strong evidence of links between the ERRFI1/EGFR axis and hallmark phenotypes of psoriatic disease - strongly suggests that the 1p36.23 risk association with psoriasis and PsA is driven by the rs11121131 enhancer and its control of *ERRFI1*. Furthermore, we have provided data which suggests a role for ERRFI1 in skin inflammation *in vivo*, further strengthening the link between psoriatic disease and ERRFI1. This is a novel gene in the pathogenesis of both psoriasis and PsA and may offer new therapeutic opportunities to target the EGFR/ERRFI1 pathway in these diseases. Inhibitors of EGFR are used in treatment of a number of cancers (30), and it is possible these drugs can be repositioned to treat PsA and Ps. Indeed, EGFR inhibitors (EGFRi) have been shown to be effective at treatment of collagen-induced arthritis in rodents (72), and psoriasis patients undergoing EGFRi treatments for cancer exhibited marked improvement in their psoriasis lesions (73). More generally, the novel association of *ERRFI1* with PsA may help broaden our understanding of its pathogenesis. Functional studies of PsA have lagged behind those of other rheumatic conditions such as RA, which may explain the comparatively poor treatment responses in this condition. It is likely that other components of this signalling pathway are also important in PsA risk, and further exploration of the EGFR/ERRFI1 axis is therefore warranted in the disease.

## Conclusions

This study has shown that rs11121131, a SNP in tight LD with the lead GWAS SNP at 1p36.23, lies in an enhancer element which controls the expression of *ERRFI1*, and that ERRFI1 is upregulated during psoriasis-like inflammation *in vivo*. In doing so, this work implicates *ERRFI1* as a novel gene in the pathogenesis of both psoriasis and psoriatic arthritis and provide a likely causal gene and variant for the 1p36.23 risk association for these conditions. By implicating *ERRFI1*, this study provides further evidence for the importance of the EGFR signalling pathway in psoriasis (62, 73), and also for the first time associates this pathway with psoriatic arthritis. This work may broaden our understanding of the pathogenesis of psoriatic disease, providing a starting point for further investigation of the ERRFI1/EGFR signalling axis in psoriasis and PsA. These investigations may facilitate the development of new treatments targeting the EGFR/ERRFI1 signalling pathway or enable drug repositioning of current EGFR inhibitors. Finally, the methods described herein are also broadly applicable to the functional analysis of other non-coding risk loci, for psoriatic disease or other common diseases.

## Methods

### Bioinformatic datasets

For general genome visualisation, UCSC genome browser (74) was used. ENCODE data (31), SNP data from dbSNP (75) and JASPAR TF predictions (33) were visualised using the integrated tracks on the UCSC browser on genome build GRCh38/hg38.

EpiMap datasets were accessed via the EpiMap repository web portal (76) and visualised on the UCSC browser on genome build GRCh37/hg19.

For determining variants in LD with other lead variants, LDProxy was used (77), typically using the European populations on genome build GRCh38/hg38.

Previous promoter capture Hi-C (35) and H3K27Ac HiChIP (44) were mapped onto genome build GRCh37/hg19 and visualised on the WashU epigenome browser (78). For details on samples used, library preparation and data analysis please refer to the relevant paper for each dataset.

### Plasmids

3^rd^ generation lentiviral packaging plasmids were obtained from Addgene. Plasmids used were: pMD2.G (Addgene #12259), pMDLg/pRRE (Addgene #12251), pRSV-REV (Addgene #12253).

HaCat dCas9-VPR cells were produced using Lenti-EF1a-dCas9-VPR-puro (Addgene #99373). HaCat dCas9-KRAB-MeCP2 cells were produced using a custom pLV-CBh-dCas9-KRAB-MeCP2- T2A-Puro synthesised by VectorBuilder.

The sgRNA delivery plasmid pLKO5.sgRNA.NeoR was cloned as follows. Briefly, the neomycin resistance gene (NeoR) was amplified from pcDNA-dCas9-p300 Core (addgene #61357) via PCR with Phusion Phusion High-Fidelity DNA Polymerase (NEB), using primers to add a 5’ BamHI site and Kozak sequence and a 3’ MluI site (For primer sequences see Additional file 2: Table S2). The PCR product was digested with BamHI-HF and MluI-HF (NEB), and purified using a Purelink quick PCR purification kit (Invitrogen). pLKO5.sgRNA.EFS.GFP (addgene #57822) was also digested with BamHI-HF and MluI-HF to remove the GFP sequence, and the backbone was isolated using QIAquick Gel Extraction kit (Qiagen). The NeoR cassette was ligated into the pLKO5 backbone using T4 DNA ligase (NEB) as per the manufacturer’s protocol. Ligated plasmid was transformed into Stbl3 E. coli (NEB) and plasmids were isolated using an E.Z.N.A endo-free plasmid mini kit II (Omega Bio-tek).

For the rs111_HDR and rs754_HDR homology donor plasmids, pUC57 (Genscript) was linearised using EcoRV-HF (NEB). Homology arms were amplified from HaCat cell genomic DNA via PCR using Q5 DNA polymerase (NEB). The −loxP-PGK-Em7-NeoR-loxP- cassettes were amplified from PL452 (79). All primers contained 30bp homologous regions to allow assembly of PCR fragments into the pUC57 backbone using 2X HiFi assembly mix (NEB) as per the manufacturer’s protocol, and ligated plasmids were transformed into Stbl3 *E. coli* cells. Confirmed plasmids were prepped for transfection using a PureLink HiPure plasmid maxiprep kit (Invitrogen). For primer sequences, see Additional file 2: Table S2.

### Cell lines and cell culture

All cells were maintained at 37°C, 5% CO2. HaCat cells were obtained from Catlag Medsystems. The HaCat dCas9-p300 stable cell line was produced from these cells as part of a previous study (35). The HaCat dCas9-VPR and dCas9-KRAB-MeCP2 stable cell lines were kindly produced by the University of Manchester Genome Editing Unit. HaCat cells were cultured in complete HaCat media (DMEM high glucose with L-Glutamine, sodium pyruvate and sodium bicarbonate (Sigma) plus 10% heat-inactivated FBS (Gibco), 1% penicillin/streptomycin (Sigma)). HaCat stable cell lines expressing dCas9 machinery were additionally cultured in 0.5µg/mL puromycin (Sigma P8833) to maintain dCas9 expression.

LentiX HEK293T cells (Clontech, catalogue #632180) were maintained in DMEM high glucose with L-Glutamine and sodium bicarbonate, without sodium pyruvate (Sigma) plus 10% heat-inactivated FBS (Gibco), 1% penicillin/streptomycin (Sigma).

### Lentivirus production

10^7^ LentiX HEK293T cells were seeded into a 15cm^2^ cell culture dish (Corning) in complete DMEM without penicillin/streptomycin and incubated overnight at 37°C, 5% CO2. Media was exchanged prior to transfection. A transfection mix was assembled in 2mL DMEM without phenol red (Gibco) as follows: 6µg transfer plasmid vector (variable), 1.5µg pMD2.G, 1.5µg pMDLg/pRRE, 3µg pRSV-Rev. Polyethylenimine (PEI) (Polyscience 23966-2) was added at a 3:1 PEI:DNA ratio and the mixture briefly vortexed and incubated at RT for 10-20 mins. The transfection mix was added to the HEK293T cells and cells were cultured for 72hrs. Viral supernatant was collected, spun at 1200RPM for 5 minutes at 4°C, and then filtered through 0.45μm cellulose acetate vacuum filter (Corning). The viral supernatant was then aliquoted and stored at −80°C.

### RNA extraction

Cells for RNA extraction were lysed in Qiazol reagent (Qiagen) and temporarily stored at - 20°C. RNA was initially extracted following the Qiazol manufacturer’s protocol using the optional 2mL MaXtract High-density gel tubes (Qiagen) to obtain the crude aqueous fraction. This was then cleaned up using a RNeasy mini RNA extraction kit (Qiagen) as per the manufacturer’s protocol, including the optional DNase treatment step.

### Guide RNA design and production

The following sgRNAs were taken from the literature: *SLC4A1* control (35), *RAB1A* control (80), Scramble2 non-targeting control (52). For sequences, see Additional file 2: Table S1.

All other sgRNAs were designed against the GRCh38/hg38 human reference genome, using both the Sanger WGE CRISPR finder (81) and CRISPOR (82). sgRNAs were designed using a balance between off-target profiles and location, and all sgRNAs with predicted 1- or 2-base mismatch off-targets or those that overlapped common variation were discounted.

For gRNAs to be delivered as Cas9 RNPs, synthetic crRNAs were synthesised by IDT, and complexed with AltR tracrRNA (IDT) *in vitro*. For lentiviral delivery, forward and reverse oligos (Sigma) were designed and annealed to produce the crRNA sequence, along with overhangs for cloning into the BsmBI site in pLKO5.sgRNA.NeoR.

### Gene expression RT-qPCR

Gene expression analysis were carried out using a TaqMan RNA-to-Ct 1-step kit (Applied Biosystems) in 384-well plate format (10µL reactions, 20ng of RNA per reaction) as per the manufacturer’s protocol, and analysed on a Quantstudio Flex 12K real-time PCR system (Applied Biosystems). All assays were carried out in triplicate. For TaqMan assay IDs see Additional file 2: Table S4.

Fold change analysis was carried out via the 2^−ΔΔCt^ method (83), using a normalisation factor calculated from the geometric mean of the Ct values of housekeeping genes *TBP* and *YWHAZ*, which better controls for outliers and variation between genes (84). For the epidermal growth factor time course, inverse ΔCt was calculated as −1*(ΔCt).

### CRISPR activation and interference in HaCat cells

For each sgRNA, the HaCat stable cell line of choice (HaCat-dCas9-VPR, HaCat-dCas9-KM2 or HaCat-dCas9-p300) was plated in 6-well TC-treated plates (Corning) at 3×10^5^ cells/well in 3mL complete HaCat media + 0.5μg/mL puromycin in triplicate, and cultured overnight. Cells for use as untransduced controls were also plated at this stage. Cells were then washed in Dulbecco’s phosphate buffered saline (DPBS) (Sigma) and media was exchanged for 2mL antibiotic-free HaCat media. 24ug polybrene (Sigma H9268) and 1mL viral supernatant containing lentivirus carrying pLKO5.sgRNA.NeoR expressing the chosen sgRNA. After 24hrs, cells were washed and media was exchanged for 3mL complete HaCat media. Cells were cultured for another 24-48hrs until wells were confluent, and then transferred to T-75 flasks in complete HaCat media and incubated overnight. Media was then exchanged for complete HaCat media + 800µg/mL Geneticin (Gibco). Selection was carried out for ∼7 days, with media being changed every 2-3 days, until all untransduced control cells had died. Cells were then moved to a maintenance concentration of Geneticin (400µg/mL) and rested overnight before being processed for RNA. sgRNA sequences can be found in Additional file 2: Table S1.

### Creation of HaCat rs11121131 and rs7548511 deletion cell lines

AltR synthetic crRNAs and AltR tracrRNA (IDT) were pre-complexed at equimolar ratios at 95°C for 5 mins and allowed to cool to RT. To assemble Cas9 RNP complexes, 0.5µL of 44µM crRNA:tracrRNA complex and 0.9µL 20µM EnGen SpyCas9 NLS (NEB) per electroporation were mixed and incubated at RT for 10-20 mins. HaCat cells were harvested and resuspended in Electroporation buffer R at a concentration of 1×10^5^ cells per 7.6µL. The electroporation mix was assembled as follows: 0.75µL 5’ gRNA RNP, 0.75µL 3’ gRNA RNP, 7.6µL HaCat cell suspension, 2µL 10.8µM AltR electroporation enhancer (IDT), 1µL 1µg/µL HDR donor plasmid (rs111_HDR or rs754_HDR). For deletion of rs11121131, rs11121131_5’_sgRNA and rs11121131_3’_sgRNA were used, and for deletion of rs7548511 rs7548511_5’_sgRNA and rs7548511_3’_sgRNA were used (For sequences, see Additional file 2: Table S1). Cells were electroporated Using the Neon transfection system 10µL kit (Invitrogen) and transferred to 0.5mL complete HaCat media + 10µM XL413 (Tocris Bioscience), alongside unedited controls. After 24hrs media was changed for complete HaCat media without XL413 and cells were cultured for 1 week. Edited cells were selected with 800μg/mL geneticin until all unedited cells had died, and were then transferred to 400µg/mL for maintenance and expansion.

To remove the −loxP-PGK-Em7-NeoR-loxP- selection cassette and establish single-cell clones, Geneticin-selected deletion cells were plated in 6-well plates at 7×10^5^ cells/well in complete HaCat media and incubated overnight. Cells were washed and media was exchanged for 1.5mL Opti-MEM (Gibco) + 5% heat-inactivated FBS (Gibco) containing AAV2-serotyped pAAV-EF1a-Cre:T2A:mNeonGreen:WPRE AAV vector (Custom, VectorBuilder) at an MOI of 10^4^ vg per cell. After 18-24hrs, media was exchanged for complete HaCat media. 48hrs post-transduction, cells were treated with TrypLE select (10X) (Gibco) for 15mins. Cells were collected and washed 3X in DPBS + 5mM EDTA, and then passed through a 40µm cell strainer (Greiner Bio-One). Cells were stained with 1µg/mL Hoechst 33342 (BD Pharmingen) at 37°C for 30 mins before being resuspended in sort buffer (PBS + 2.5mM EDTA + 25mM HEPES (Sigma-Aldrich)). Sorting was carried out by the University of Manchester Flow Cytometry Core Facility on a BD Influx cell sorter to isolate mNeonGreen^+^ Hoechst 33342^Low^ single cells. Individual cells were sorted into each well of a 96-well flat bottom plate (Corning) in 100uL of complete HaCat media. Single cell clones were monitored, expanded, and utilised for downstream experiments.

Note that prior to creating the deletion lines, sgRNAs were assessed for on-target efficiency via Synthego’s ICE analysis tool. Briefly, HaCat keratinocytes edited with individual sgRNAs delivered as Cas9 RNPs via electroporation with the Neon transfection system as previously described, and harvested 72hrs post-editing. Genomic DNA was isolated using PureLink Genomic DNA mini kit (Invitrogen) as per the manufacturer’s protocol. PCRs were then carried out as described previously to amplify a 300-600bp region in which the Cas9 cut site was approximately central. These products were then sent for Sanger sequencing by the University of Manchester Genomic Technologies Core Facility, using the forward primer used for the PCR as the sequencing primer. Resulting sequencing data was used to determine editing efficiency using the ICE analysis web tool (85).

### Epidermal Growth Factor stimulation time course

The EGF stimulation time course was carried out across four time points: 0hr, 2hr, 4hr and 24hr. For each time point, 3 clones of rs11121131 deletion HaCat cells, 3 clones of rs7548511 deletion HaCat cells were seeded in 24-well plates at 1.5×10^5^ cells/well in complete HaCat media. For the control, 3 wells of polyclonal normal HaCat cells were seeded for each timepoint. Cells were incubated overnight, and then at time 0 media was exchanged for complete HaCat media + 20ng/mL human recombinant EGF (Gibco PHG0311L) for all wells except the 0hr timepoint, which were processed immediately. At each respective timepoint, cells were washed with DPBS and lysed directly in Qiazol reagent using a sterile miniature cell scraper (Biotium). Cell lysate was collected and stored for subsequent RNA processing and analysis.

### ChIP-qPCR

Fixation and chromatin immunoprecipitation was carried out according using an iDeal ChIP-qPCR kit according to the manufacturer’s protocol, with some modifications.

A confluent T-75 flask of HaCat cells (∼10^7^ cells) was fixed in 1% formaldehyde for 10 mins, before being quenched, harvested and processed for chromatin shearing according to the manufacturer’s protocol. For shearing, the nuclei pellet was resuspended in 1mL of shearing buffer + protease inhibitor (Diagenode), incubated for 10 mins on ice, and then transferred to a 1mL 12×12mm milliTUBE (Covaris) for sonication. Chromatin was sonicated using the Covaris S220 focused ultrasonicator (20 mins, PIP = 120W, duty factor 5%, 200 cycles/burst, 4-6°C), and then diluted to the equivalent of 4.2×10^5^ cells per 100μL. Debris was removed by centrifugation at 16000xg for 10 mins at 4°C, and supernatant was taken forward for immunoprecipitation.

Immunoprecipitation was carried out as per the kit protocol, using using either 2μg Anti-Histone H3 (acetyl K27) antibody - ChIP Grade (Abcam ab4729), 7μL Tri-Methyl-Histone H3 (Lys27) (C36B11) Rabbit mAb (Cell Signalling Technology #9733) or 1μg negative control rabbit IgG (Diagenode) per ChIP reaction. 1% input chromatin samples were kept for later analysis. Final immunoprecipitated DNA samples from the kit were then purified using a Monarch PCR and DNA cleanup kit (5μg) (NEB), with a final elution volume of 50μL in TE buffer. Immunoprecipitated DNA and input samples were stored at −20°C until needed.

qPCR analysis of immunoprecipitated DNA and input controls was carried out using using SsoAdvanced Universal SYBR Green Supermix (Bio-Rad) and the QuantStudio 12K Flex RT-PCR instrument (Applied Biosystems) in 384-well plate format (10µL reactions) in triplicate. qPCR primers (Sigma) were designed to amplify 75-200bp amplicons and were checked for specificity using Primer-BLAST (ncbi.nlm.nih.gov), alongside melt curve analysis and gele electrophoresis to check for single products. Primer efficiency was calculated using a 10-fold serial dilution of Human Mixed Genomic DNA (Promega G3041). For primer sequences and calculated efficiencies, see Additional file 2: Table S3. Average Ct values were plotted against log(gDNA concentration), followed by line fitting and slope analysis (Efficiency (%) = (10^(−1/slope)^−1) x 100) to select primers with 90 – 110% efficiency. Primers were used at 250nM. The following commercial control qPCR assays were also used: GAPDH positive control ChIP primer pair (Abcam ab267832), Human Positive Control Primer Set MYT1 (Active Motif, #71007), Human Negative Control Primer Set 1 (Active Motif, #71001), Human Negative Control Primer Set 3 (Active Motif, #71023). For analysis, 1µL of either immunoprecipitated DNA or input control DNA was used per reaction. % recovery of chromatin relative to input controls was calculated as per the kit protocol using the following equation: % recovery = _2_((Ct(1% input) – 6.64) - Ct(Sample)) _x 100._

### IF staining of mouse skin samples

All animal experiments were locally ethically approved and performed in accordance with the UK Home Office Animals (Scientific Procedures) Act 1986 under PPL: PP7875093. Eight week old male C57BL/6 mice (Charles River) were treated topically on their ears with 10 mg Aldara cream (Meda Pharmaceuticals, containing 5% Imiquimod and other inflammatory constituents) per ear, daily for 3 days to induce inflammation. Untreated mice were used as control group. The day following the final treatment, mice were euthanised and ear skin was frozen in OCT compound (Tissue Tek). Ear tissues were sectioned (7µm) using a cryostat, and were stored at −20°C. For immunofluorescence staining, tissues sections were thawed on ice for 30 min, rinsed in PBS for 10 min at RT and blocked using blocking buffer (Fish Gelatin blocking agent, BIOTIUM) in the dark for 1 hr. MIG-6 ERRFI1 Rabbit polyclonal antibody (Protein tech) was used (diluted as per manufacturer’s instructions) for 2 hr. Alexa fluor 488 goat anti rabbit IgG (H+L) (Invitrogen) was used as per manufacturer’s instruction. CoraLite 555-cojugated cytokeratin 19 polyclonal antibody (Proteintech) was used as cytokeratin marker. Recombinant rabbit IgG (Abcam) and CoraLite 555-cojugated rabbit IgG control polyclonal antibody (Proteintech) were used as isotype controls for the experiments at the same concentrations. Tissue sections were then incubated for 5 min with Ready Probes Tissue Autofluorescence Quenching Kit (Invitrogen). Slides were mounted with VECTASHIELD HardSet Antifade Mounting Medium with DAPI (Vector Laboratories, INC.), and images were acquired using a Zeiss Axioimager.D2 microscope and captured using a Coolsnap HQ2 camera (Photometrics) using MetaVue Software (Molecular Devices).

### Inflammation measures in the psoriasis-like inflammation model

The psoriasis-like skin inflammation model was carried out as described in the Immunofluorescence methods section. On each day of skin treatment, the ear skin thickness was measured by digital micrometer (Mitutoyo), and the redness and scaling were visually scored using a scale. On day 3 mice were euthanized and ear skin and draining lymph nodes (auricular) were analyzed.

### H&E staining

Skin was fixed in 10% neutral buffered formalin, embedded in paraffin and cut to 5 μm. Haematoxylin and eosin staining was performed by a Shandon Varistain V24-4. Image acquisition used a 3D-Histech Pannoramic-250 microscope slide-scanner and Case Viewer software (3D-Histech).

### Flow cytometric analysis of cells

Ears were peeled in half and digested with 0.8% w/v Trypsin (Sigma) for 30 min, then chopped and digested in 0.1 mg/ml (0.5 Wunch units/ml) Liberase TM (Roche) at 37°C for 1 h. Skin and auricular lymph node cells were passed through 70 µm cell strainers and counted using a haemocytometer and trypan blue (Invitrogen) exclusion. Cells were incubated with 0.5 µg/ml anti-CD16/32 (2.4G2, BD Bioscience), Near IR Dead cell stain (Invitrogen) and fluorescently labeled antibodies. For analysis of IL-17 production, cells were cultured for 4 h with 10 μM Brefeldin A before staining. Cells were fixed and permeabilized with Foxp3/Transcription Factor Buffer Staining Set (eBioscience) and were analyzed using a BD LSRII flow cytometer and FlowJo (TreeStar). For all antibodies used, see Additional file 2: Table S5.

### Statistics and graphs

All graphs were produced in Graphpad Prism 9.

RT-qPCR experiments were analysed via one-way ANOVA analyses with post-hoc Tukey’s tests using Graphpad Prism 9.

ChIP-qPCR data were analysed by student’s T-tests for each treatment vs controls using Graphpad Prism 9.

For the assessments of mouse ear inflammation, normality was tested by Shapiro−Wilk test. Unpaired t-tests (with Welch’s correction where required) were used to determine significant differences. Analysis was carried out using Graphpad prism 9.

Analysis of the EGF time course data by linear mixed effect model was carried out in RStudio with R version 4.0.5 and RTools40, using the nlme package (available via CRAN). The model was built using the lme() command to include the fixed effects of deletion class, time, and the interaction of deletion class and time, with sample ID as a random effect.

## Declarations

### Ethics approval and consent to participate

Experiments involving animals in this study were carried out in accordance with the UK Home Office Animals (Scientific Procedures) Act 1986 under PPL: PP7875093

### Consent for publication

Not Applicable

### Availability of data and materials

Capture Hi-C datasets from Ray-Jones *et al.,* 2020 which were analysed are available in the GEO repository under accession number GSE137906.

HiChIP datasets from Shi *et al*., 2021 are available in the GEO repository under accession number GSE151193.

### Authors’ contributions

OJG, AA, SE, HRJ and CS contributed significantly to the design and conception of the work herein. OJG, SR, AS and MP were involved in the acquisition and analysis of the data within this manuscript. OJG, SR, AS, AA and SE contributed to the interpretation of results obtained. All parties contributed to the drafting and revision of the manuscript.

## Acknowledgements

The authors would like to thank Antony Adamson and the University of Manchester Genome Editing Unit for their consultation and experimental assistance on this project. The authors acknowledge assistance from Peter Walker, Roger Meadows, and Gareth Howell, and the use of the University of Manchester Histology, Flow Cytometry, and Biological Services facilities. The University of Manchester Bioimaging Facility microscopes used in this study were purchased with grants from BBSRC, Wellcome, and the University of Manchester Strategic Fund. Thanks to BMC biology for allowing reproduction of a figure from their publication for figure 2A of this manuscript.

## Figures

**Figure S1:**
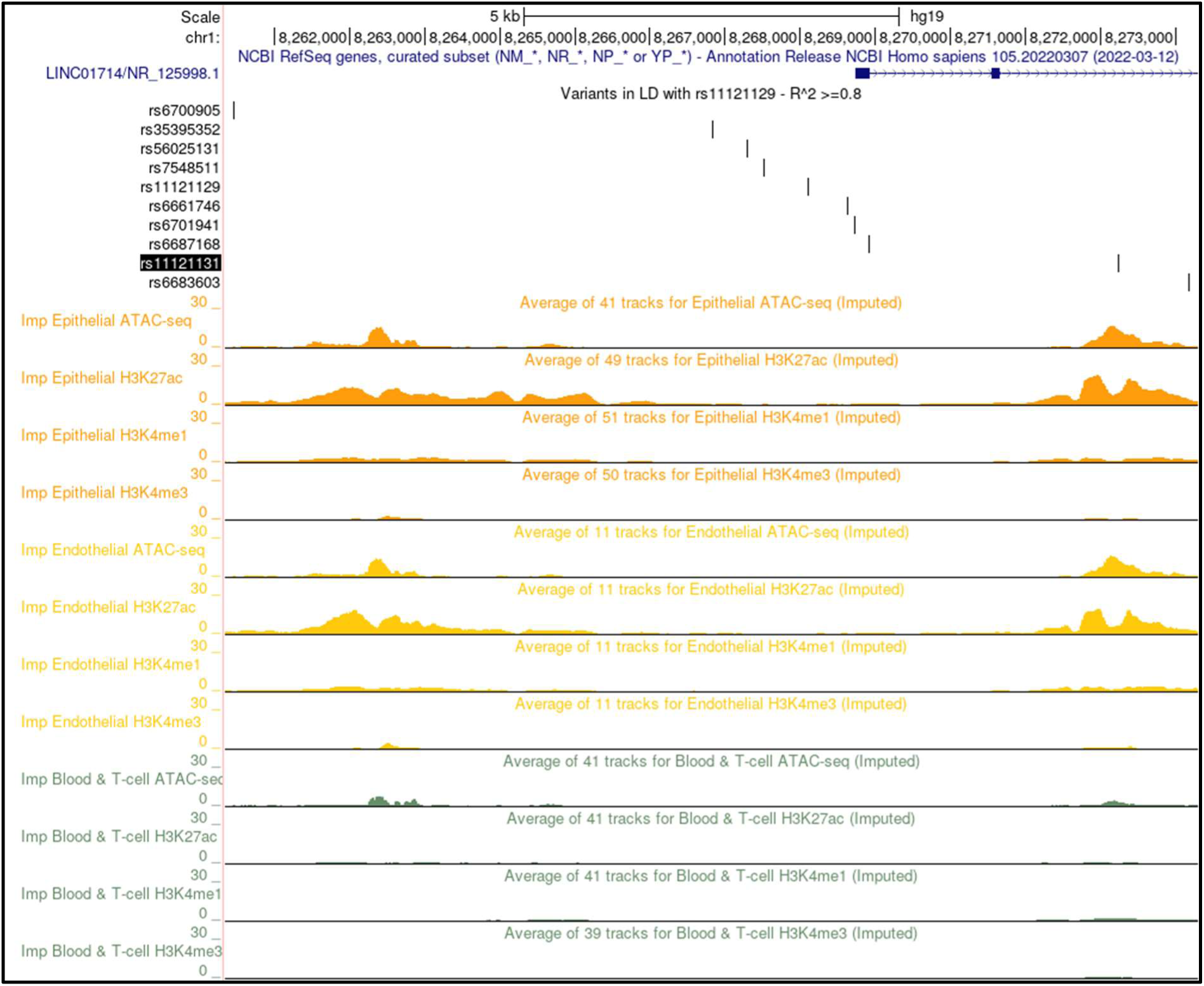
Prioritised SNP set with EpiMap average track data. UCSC image (GRCh37/hg19) showing the prioritised SNPs at 1p36.23 overlayed with data from the EpiMap dataset (32), showing average tracks for imputed ATAC-seq peaks, H3K27Ac, H3K4Me1 and H3K4Me3 histone modifications across 3 tissue categories: Epithelial, Endothelial and Blood & T-cell.

**Figure S2:**
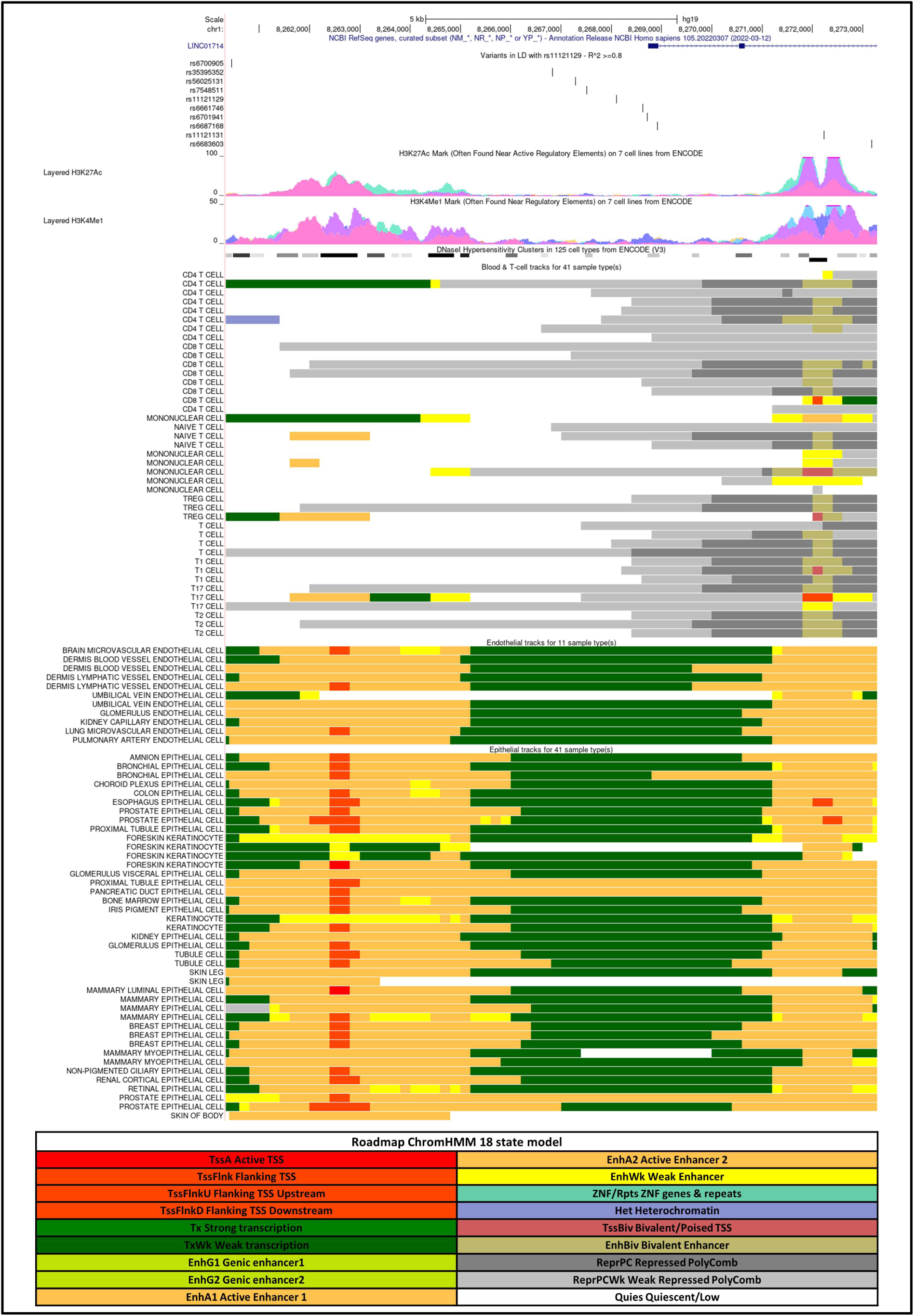
EpiMap ChromHMM: ChromHMM. analysis of samples from the EpiMap database (32) at the 1p36.23 risk locus, visualised in the UCSC genome browser (GRCh37/hg19). Data shown for tissues under the category “Blood & T-cell” and “Epithelial”. The bottom table shows the colour key for the ChromHMM model classifications.

**Figure S3:**
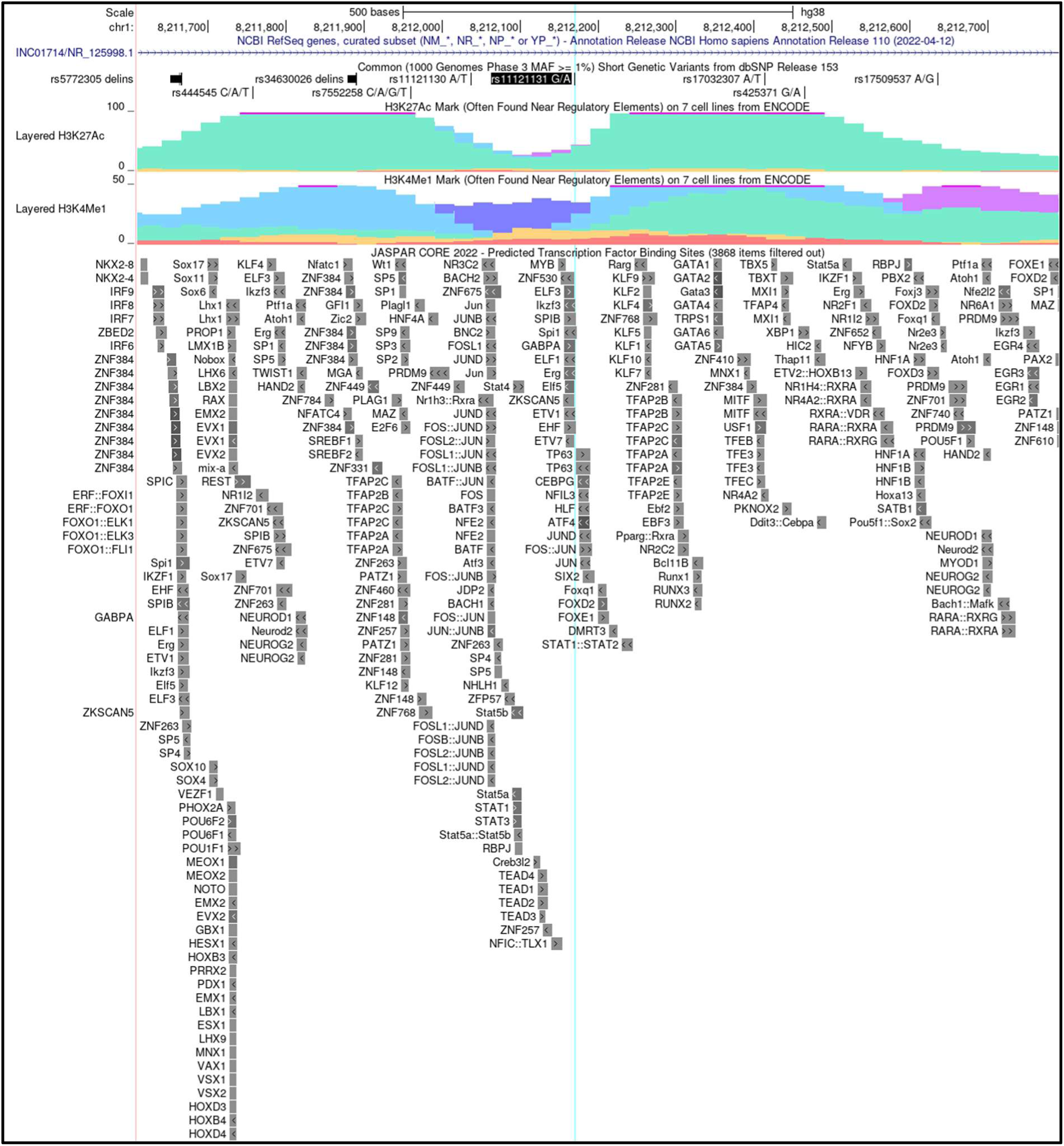
JASPAR transcription factor predictions at rs11121131. UCSC image (GRCh38/hg38) showing the region surrounding rs11121131 (vertical blue highlight) overlayed with predicted transcription factor binding sites from the JASPAR core motifs 2022 dataset (33).

**Figure S4:**
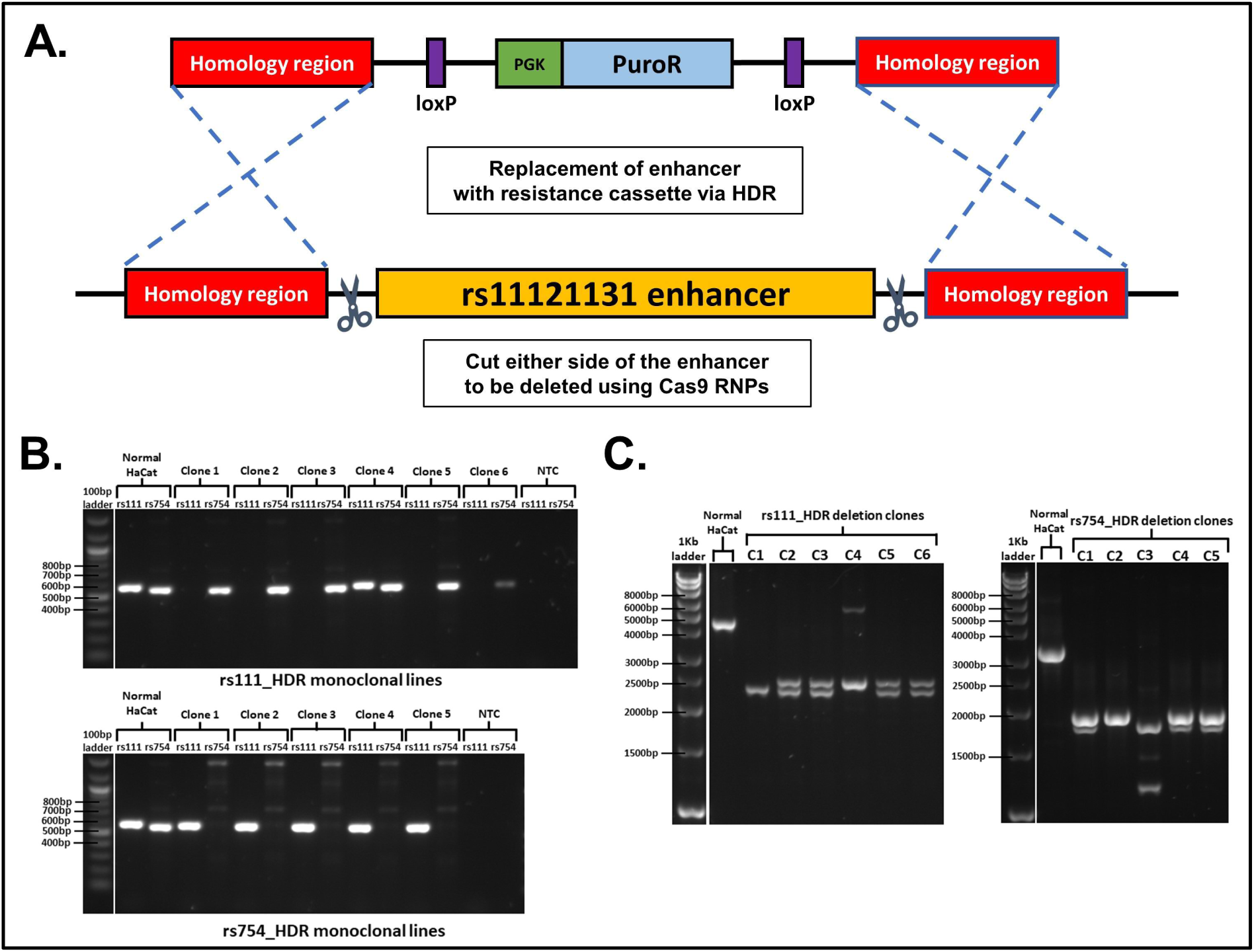
Deletion strategy and clone confirmation for rs11121131 and rs7548511 deletion clones. **A.**Schematic detailing the strategy utilised for deletion of the rs11121131 and the rs7548511 loci in HaCat cells. Cas9 RNPs targeting either side of the locus are delivered alongside a double-stranded DNA homology donor. Homologous repair leads to replacement of the locus with a Neomycin resistance marker for selection of cells harbouring a successful deletion. The resistance marker is flanked by loxP sites, allowing subsequent removal via transient expression of Cre recombinase. **B.** Gels showing internal PCR amplicons for all the rs111_HDR and rs754_HDR clones, detecting the presence of either the rs11121131 (rs111) or rs7548511 (rs754) locus. **C.** Gels showing PCR amplicons amplifying across the whole deletion site of either the rs11121131 locus or the rs7548511 locus in their respective deletion clones, utilising PCR primers outside of the HDR donor homology arms. Expected size for wild-type locus = 4449bp, expected size for HDR deletion allele with NeoR sequence successfully excised = 2477bp. For primer sequences see Additional File 2: Table S2.

**Figure S5:**
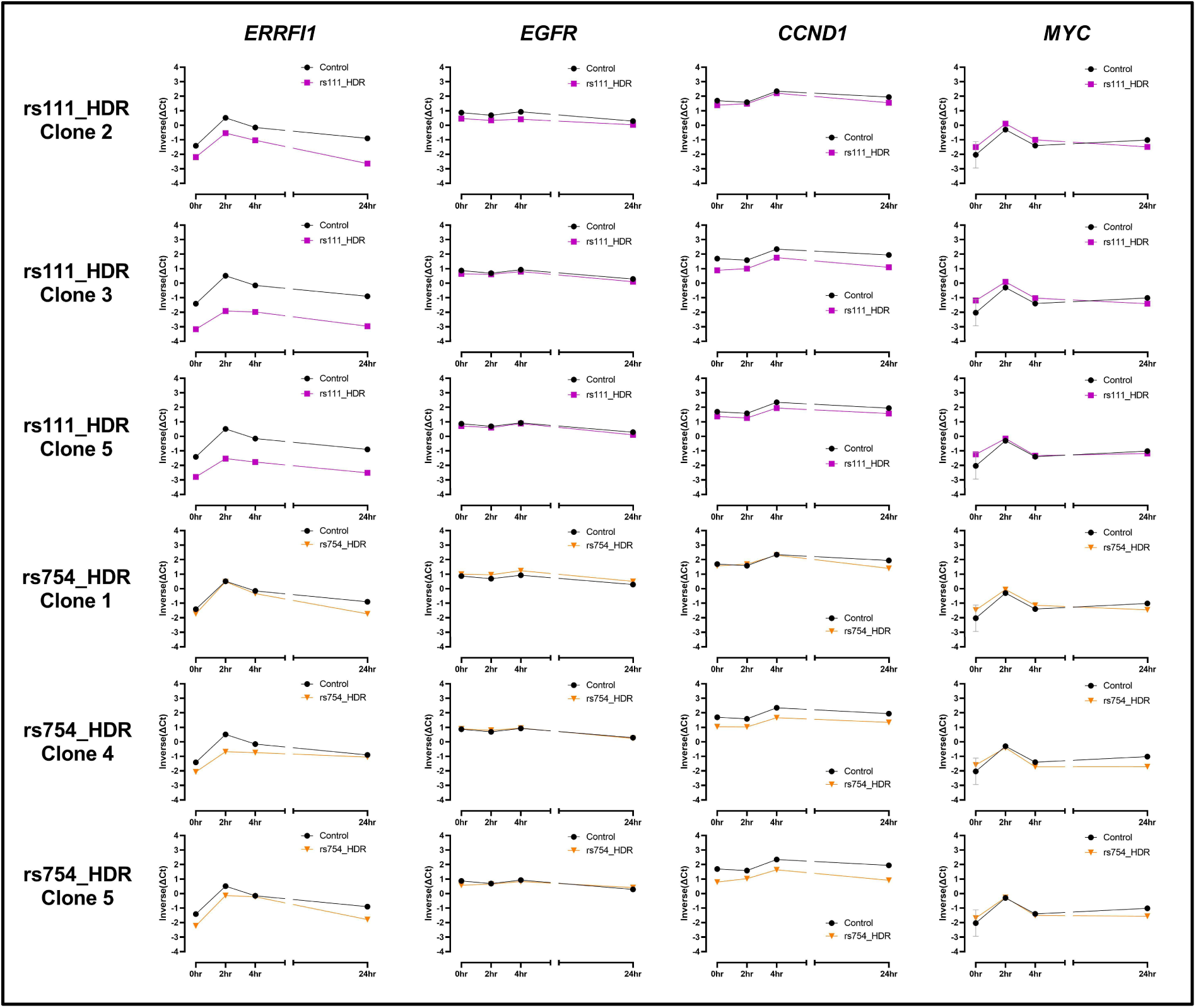
Individual clone data from EGF stimulation time course. EGF stimulation time course data from Figure 3C, with data separated into individual deletion clones. In each panel, the data from one HaCat deletion control is shown against a control line showing the mean of all 3 control samples.

**Figure S6:**
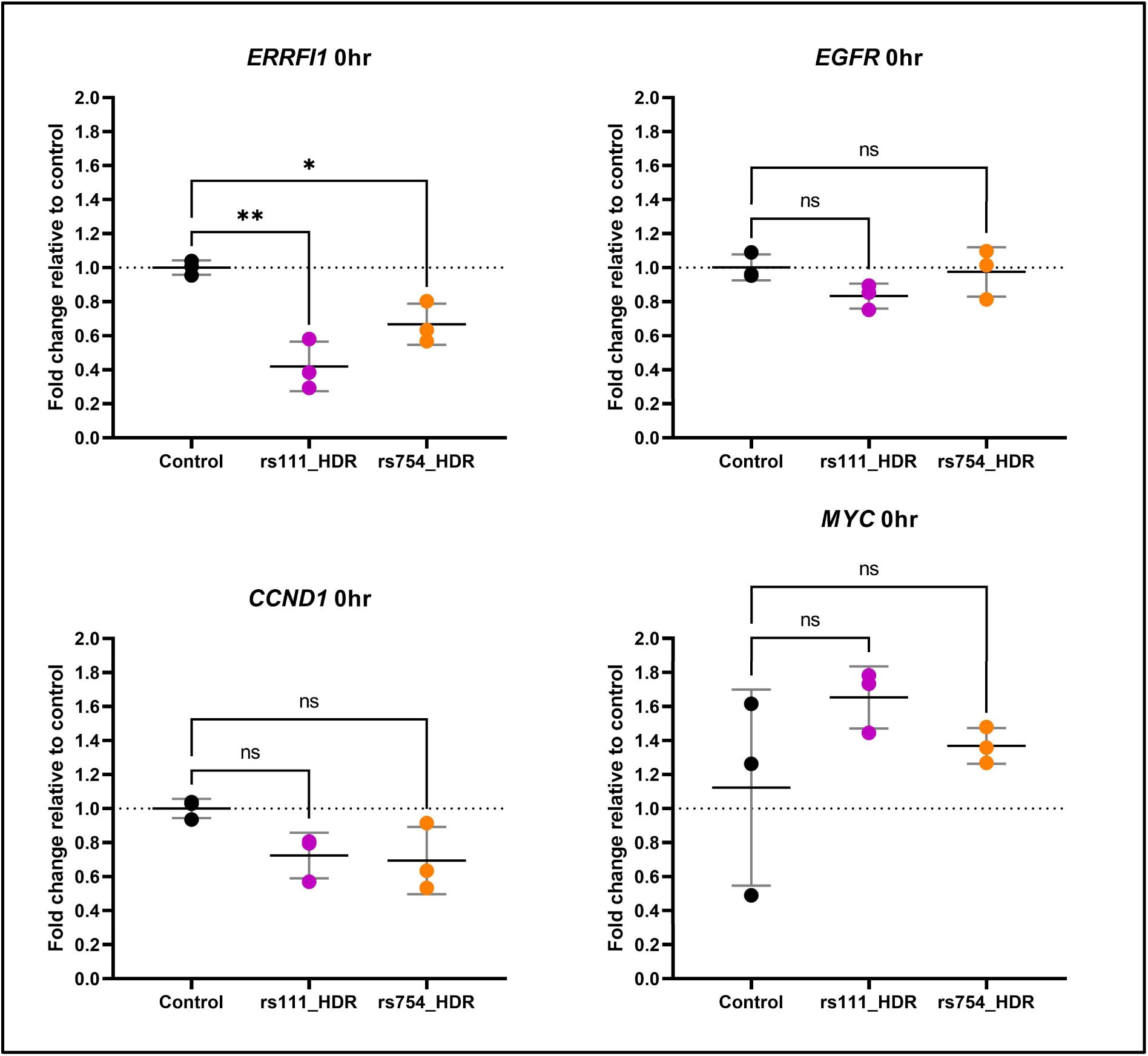
Fold change relative to control for 0hr samples from EGF time course ΔCt. Fold change in mRNA levels calculated relative to controls from the 0hr timepoint samples from the EGF timecourse. Fold change was calculated relative to the average Ct for the 3 control replicates using the 2^-ΔΔCt^ method. Dots show either individual clones for the rs111_HDR and rs754_HDR deletion samples (n=3) or the 3 individual replicates of polyclonal HaCat control cells (n=3). Data analysed via one-way ANOVA with a post-hoc Tukey’s test to determine significance relative to pLKO5.NeoR (empty) control (ns: P≥0.05; *: P<0.05; **: P<0.01; ***: P<0.001; ****: P<0.0001).

**Figure S7:**
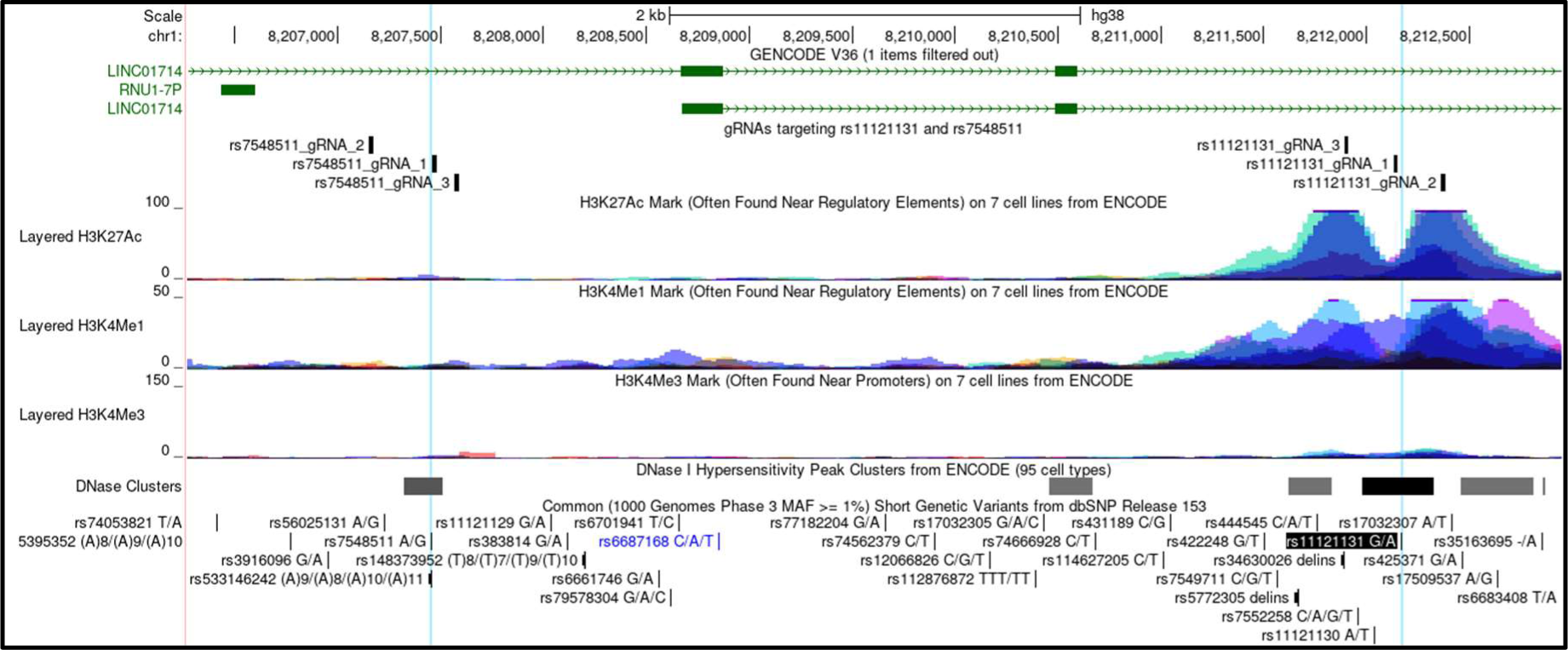
Locations of CRISPRa and CRISPRi guide RNAs at rs11121131 and rs7548511. UCSC image (GRCh38/hg38) showing locations of the sgRNAs targeting either the rs11121131 or the rs7548511 loci for the CRISPRa and CRISPRi experiments. Positions of rs11121131 and rs7548511 are shown by the vertical blue highlights.

**Figure S8:**
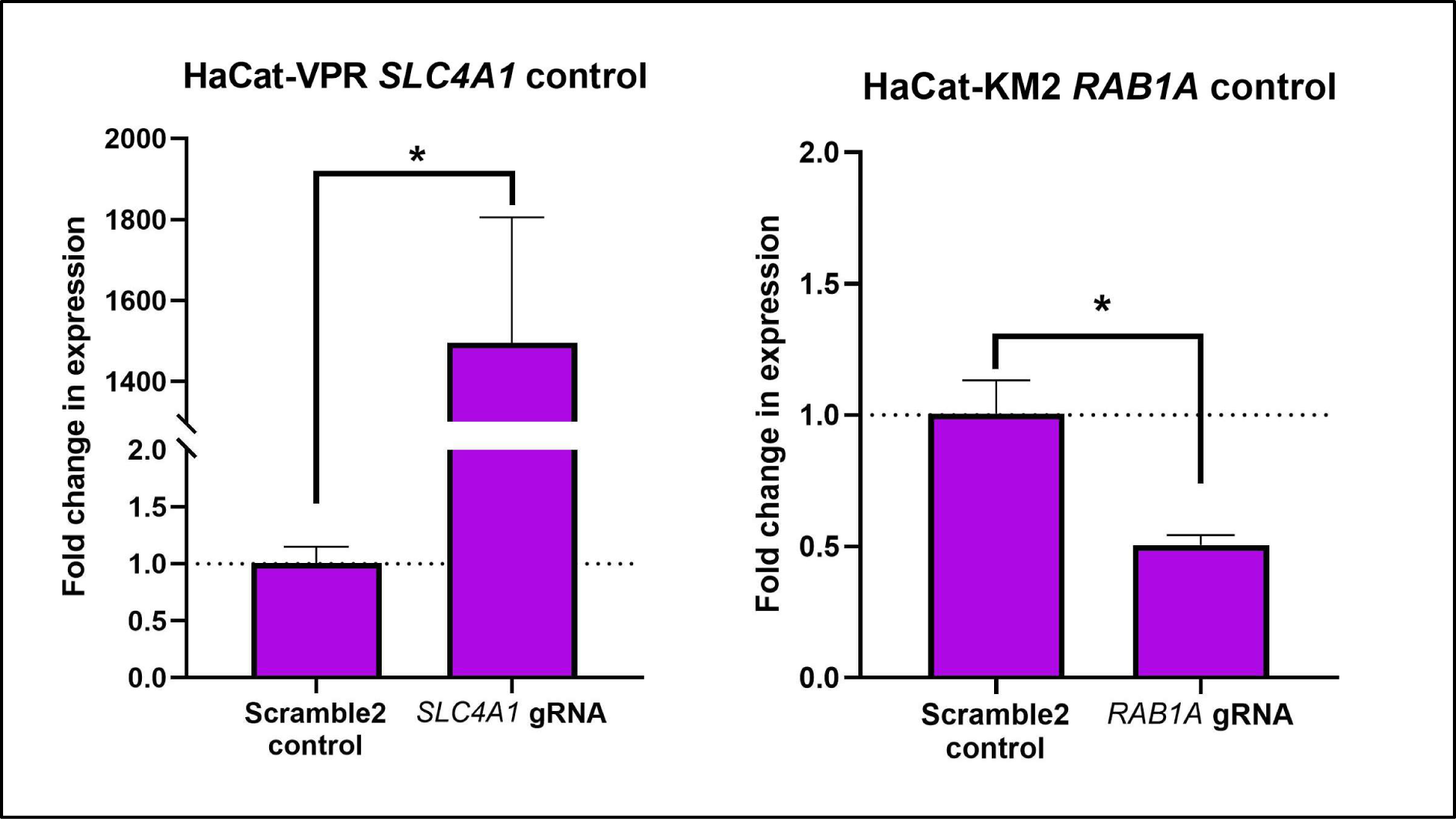
HaCat dCas9-VPR and dCas9-KRAB-MeCP2 positive control tests. Positive control tests for the HaCat-dCas9-VPR and HaCat-dCas9-KRAB-MeCP2 stable cell lines. The positive control sgRNA (targeting either SLC4A1 for activation or RAB1A for repression) was compared to Scramble2 controls. n=3 biological replicates per condition, error bars show mean with standard deviation. For RT-qPCR, 20ng RNA was used per assay, and all assays were carried out in triplicate. Fold change calculated relative to the mean of the pLKO5.NeoR (empty) controls via the 2^-ΔΔCt^ method. Where the transcript was undetectable in the control sample, the sample was assigned a Ct of 40 for analysis. Data analysed via student’s t-test to determine significance relative to Scramble2 control (ns: P≥0.05; *: P<0.05; **: P<0.01; ***: P<0.001; ****: P<0.0001).

**Figure S9:**
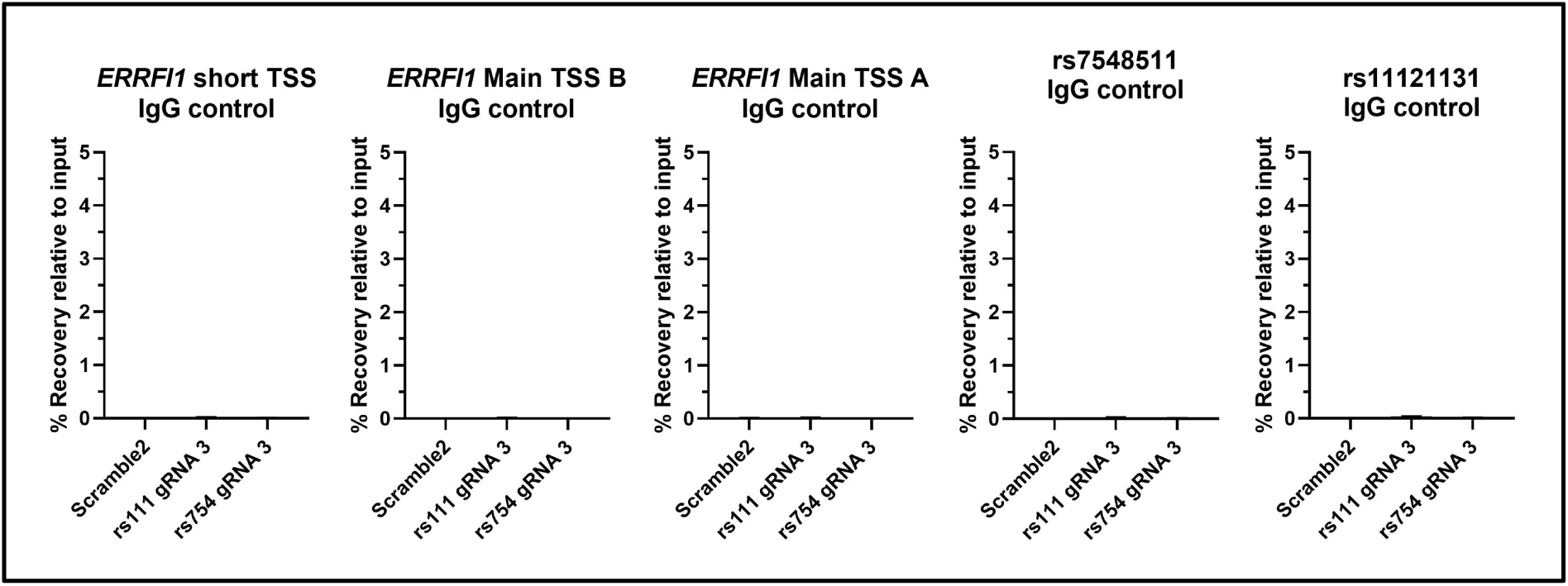
ChIP-qPCR IgG controls. IgG control immunoprecipitations from the ChIP-qPCR experiment shown in figure 6. n=2 biological replicates per condition. qPCR was carried out using 1μL of sample per 10μL reaction using the BioRad SsoAdvanced universal SYBR green Supermix, with 3 technical replicates per reaction.

**Figure S10:**
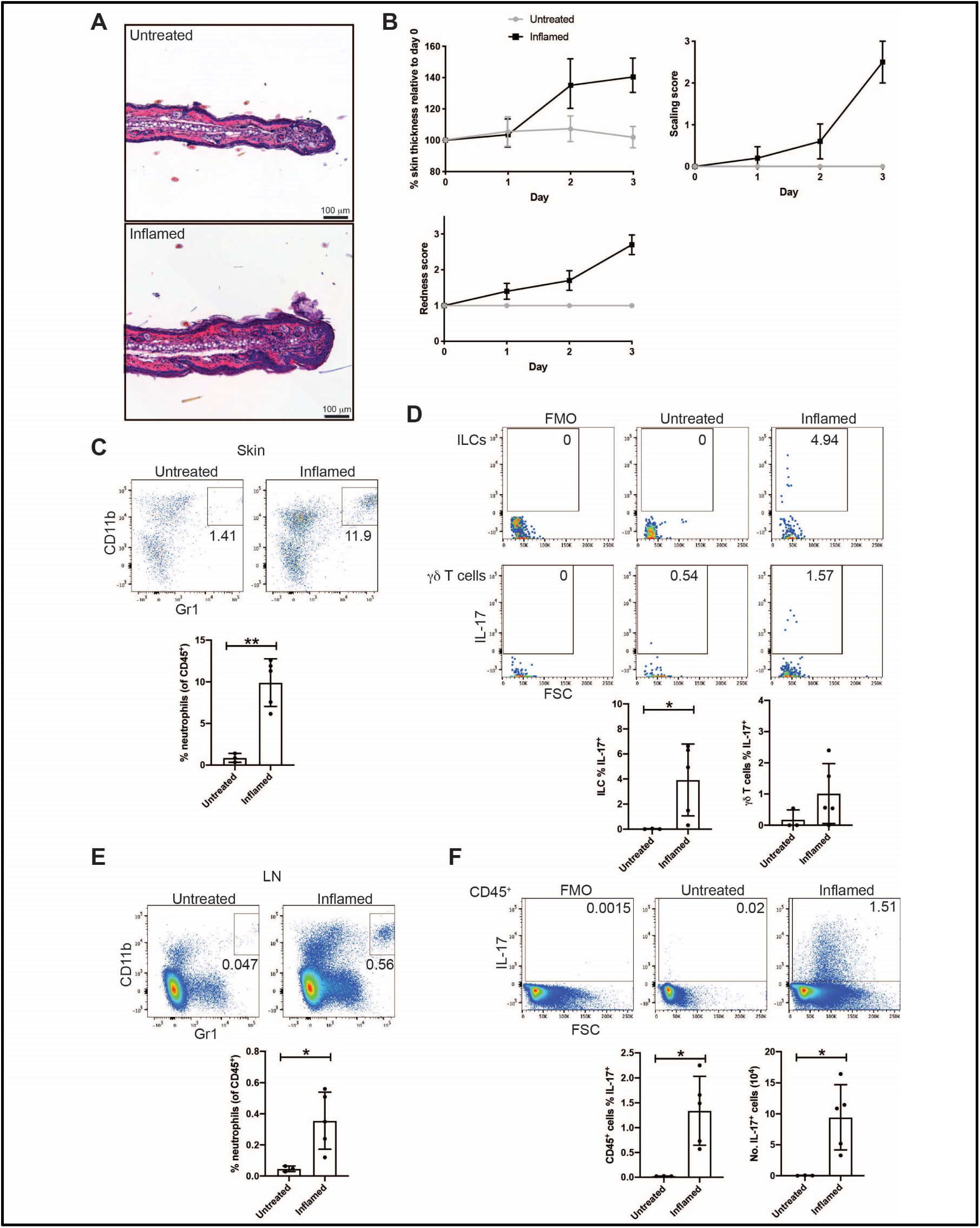
Aldara cream treatment results in psoriasis-like skin inflammation in mouse ears. C57BL/6 mice were treated with 10mg Aldara cream topically on each ear for 3 days. A day after the final treatment, mice were euthanised and ear skin and skin draining lymph nodes (auricular) were analysed for inflammation. **A.** Ear skin was sectioned and stained with haematoxylin and eosin. **B.** Ear redness and scaling was scored daily, and ear thickness was measured daily using a digital micrometer. **C-D**. Ear skin cells were isolated and analysed by flow cytometry. **C.** Neutrophils (CD11b^hi^ Gr1^hi^) were gated within the CD45^+^ cell population. **D.** IL-17 production was examined within the innate lymphoid cell (ILC) (Live CD45^+^ Lineage^-^ [CD11b CD11c Gr1 F4/80 Ter119 FceRIa CD19] CD3^-^ TCRgd^-^ TCRb^-^ Thy1^+^ CD127^+^) and dermal gd T cell (Live CD45^+^ CD3^mid^ TCRgd^mid^) populations. FMO indicates the fluorescence minus one control (stained with all antibodies except for IL-17). FSC indicates the forward scatter signal. **E-F.** Auricular lymph node cells (LN) were analysed by flow cytometry. **E.** Neutrophils (CD11b^hi^ Gr1^hi^) were gated within the CD45^+^ cell population. **F.** IL-17 production was examined within the live CD45^+^ population. Data were analysed by unpaired t test, with Welch’s correction applied to D-F. Statistically significant differences are indicated by * for p < 0.05 and ** for p < 0.01. n = 3 for uninflamed and n = 5 for inflamed.

**Figures S11.**
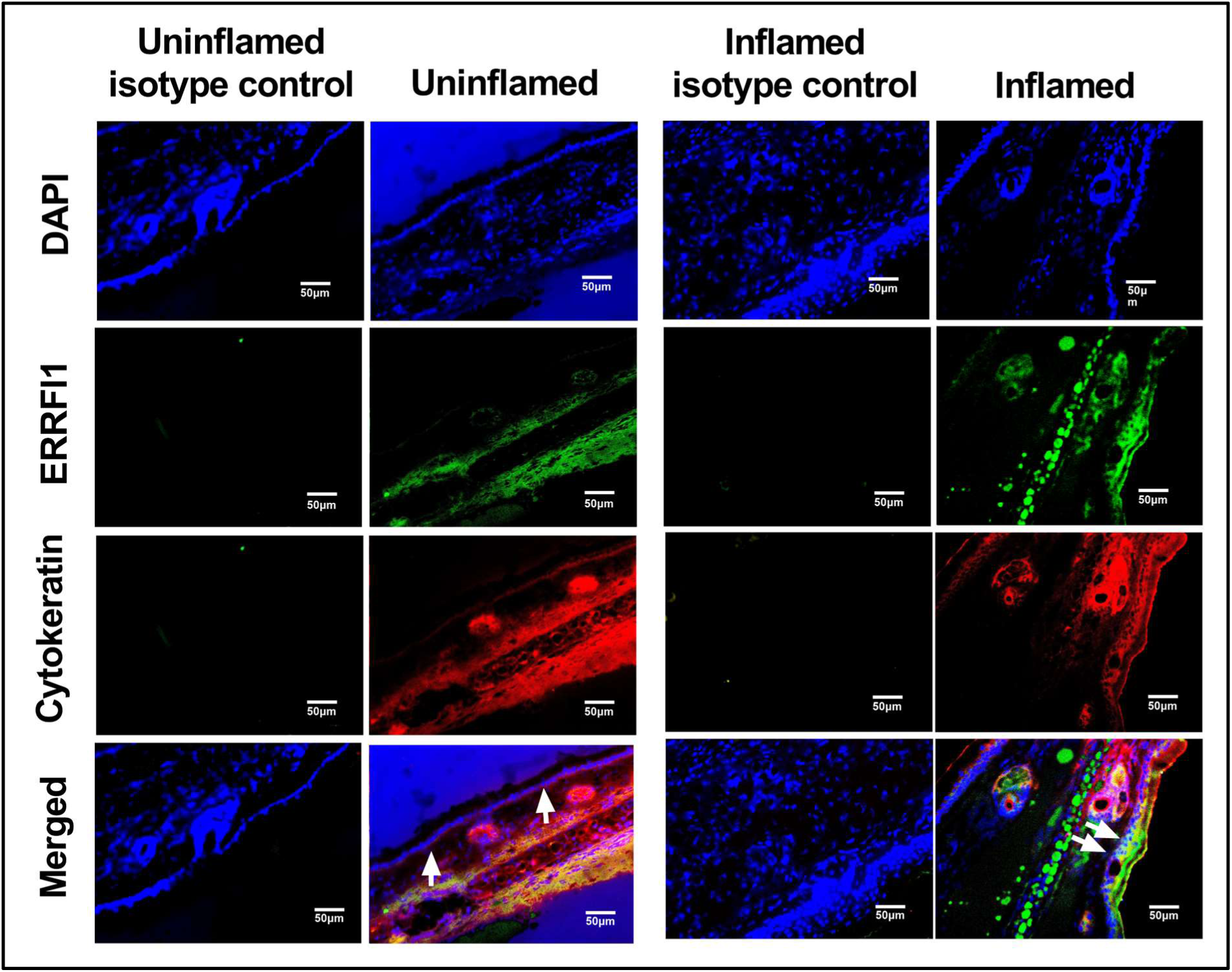
Additional murine ERRFI1 immunofluorescence images for figure 7. Figures for the additional animals (5 imiquimod-treated, 5 controls) from representative main text figure 7, showing immunofluorescent imaging of sections taken from the ear skin of imiquimod-induced psoriasis-like mouse models vs uninflamed controls. Each figure (11-15) shows the staining for one imiquimod-treated mouse and one uninflamed control mouse. C57BL/6 mice were treated topically with Imiquimod-containing Aldara cream daily for 3 days to induce psoriasis-like skin inflammation. C57BL/6 mice ear tissues were harvested, cryo- sectioned and stained for ERRFI1 expression, alongside cytokeratin 19 to help identify the epidermal keratinocyte layer and a DAPI nuclear co-stain. Controls were taken from untreated C57BL/6 mice. White pointers indicate the epidermal layer. Appropriate isotype controls were used to confirm the specificity of the observed staining. Scale bars show 50µm.

**Figures S12:**
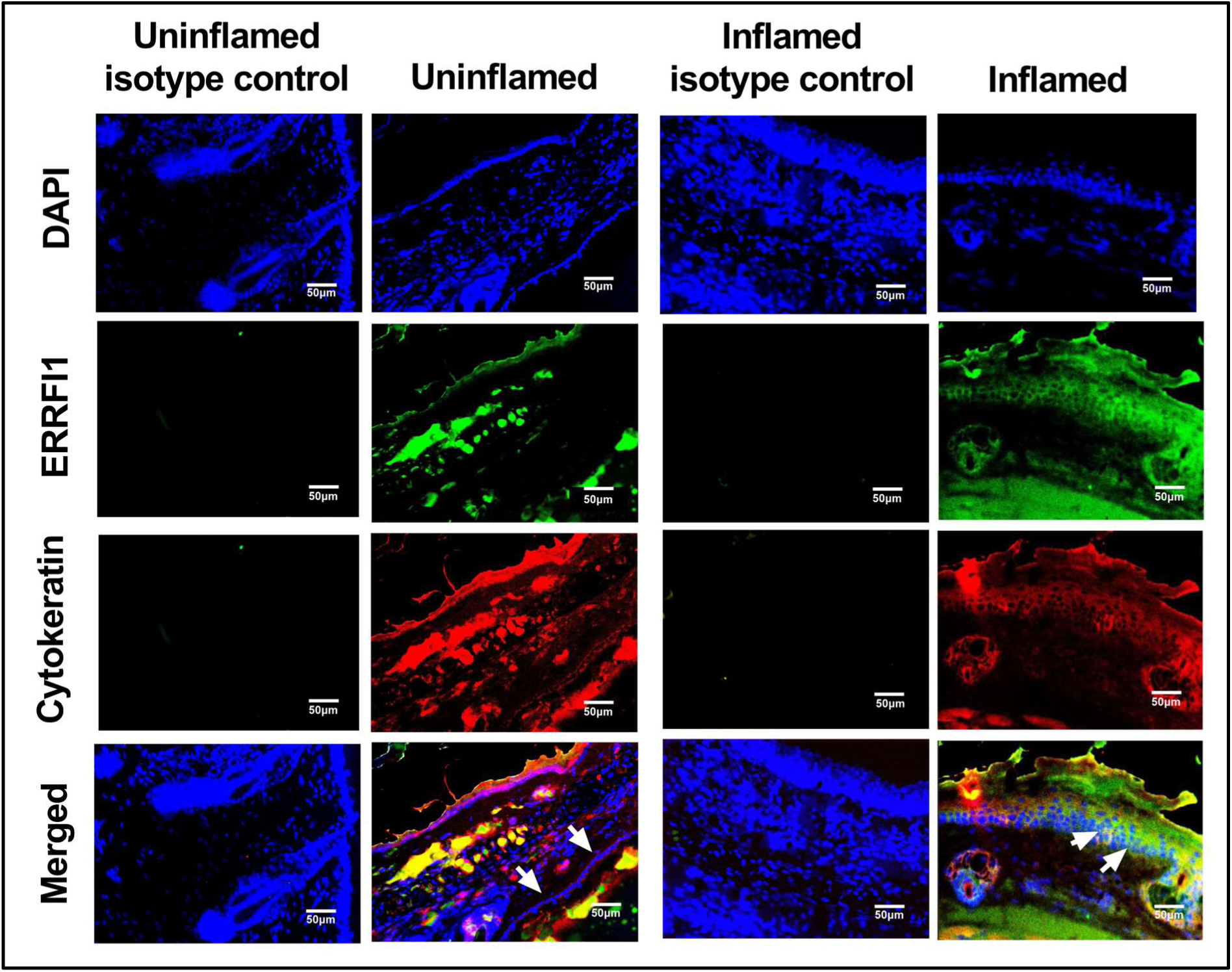
Additional murine ERRFI1 immunofluorescence staining 2.

**Figures S13:**
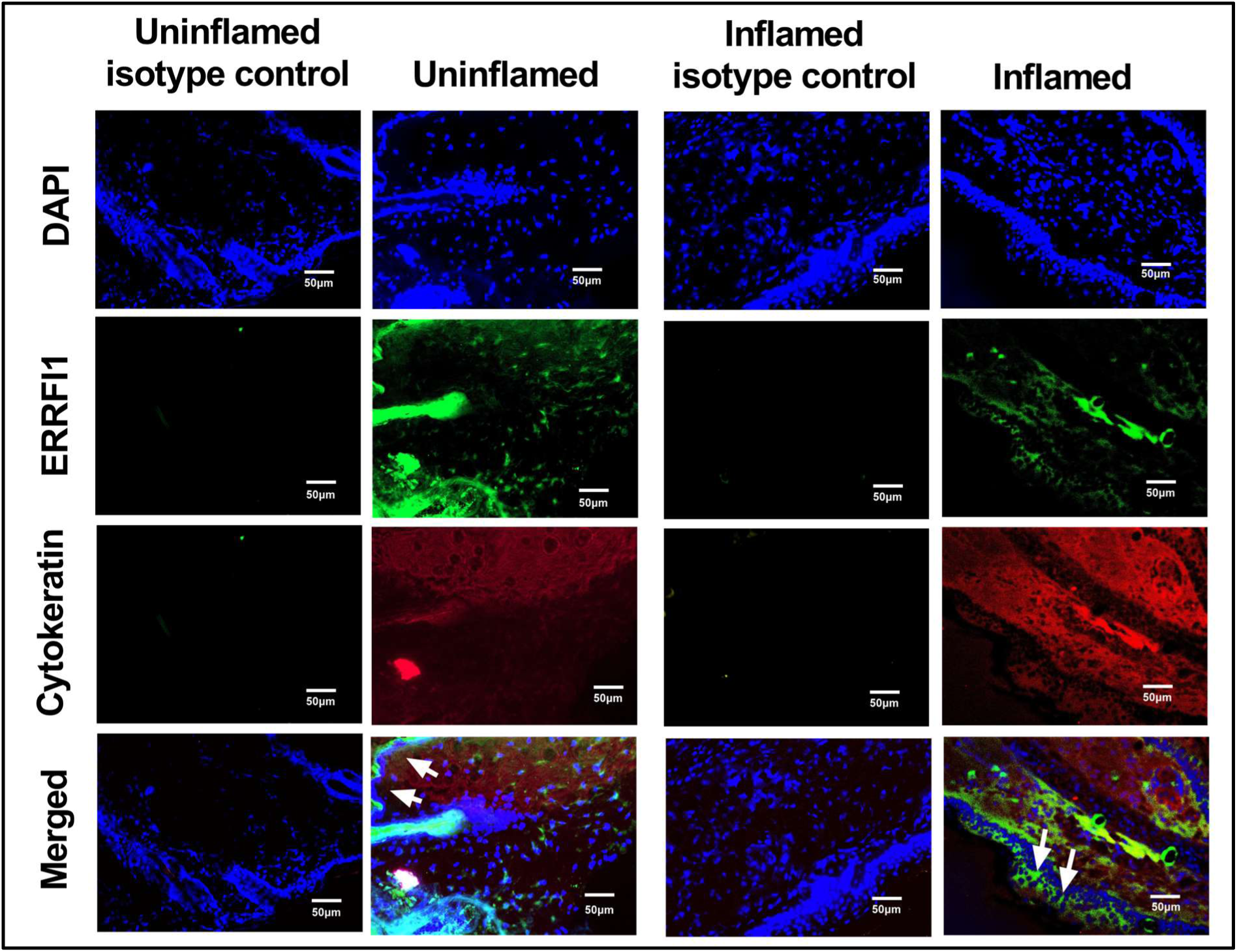
Additional murine ERRFI1 immunofluorescence staining 3.

**Figures S14:**
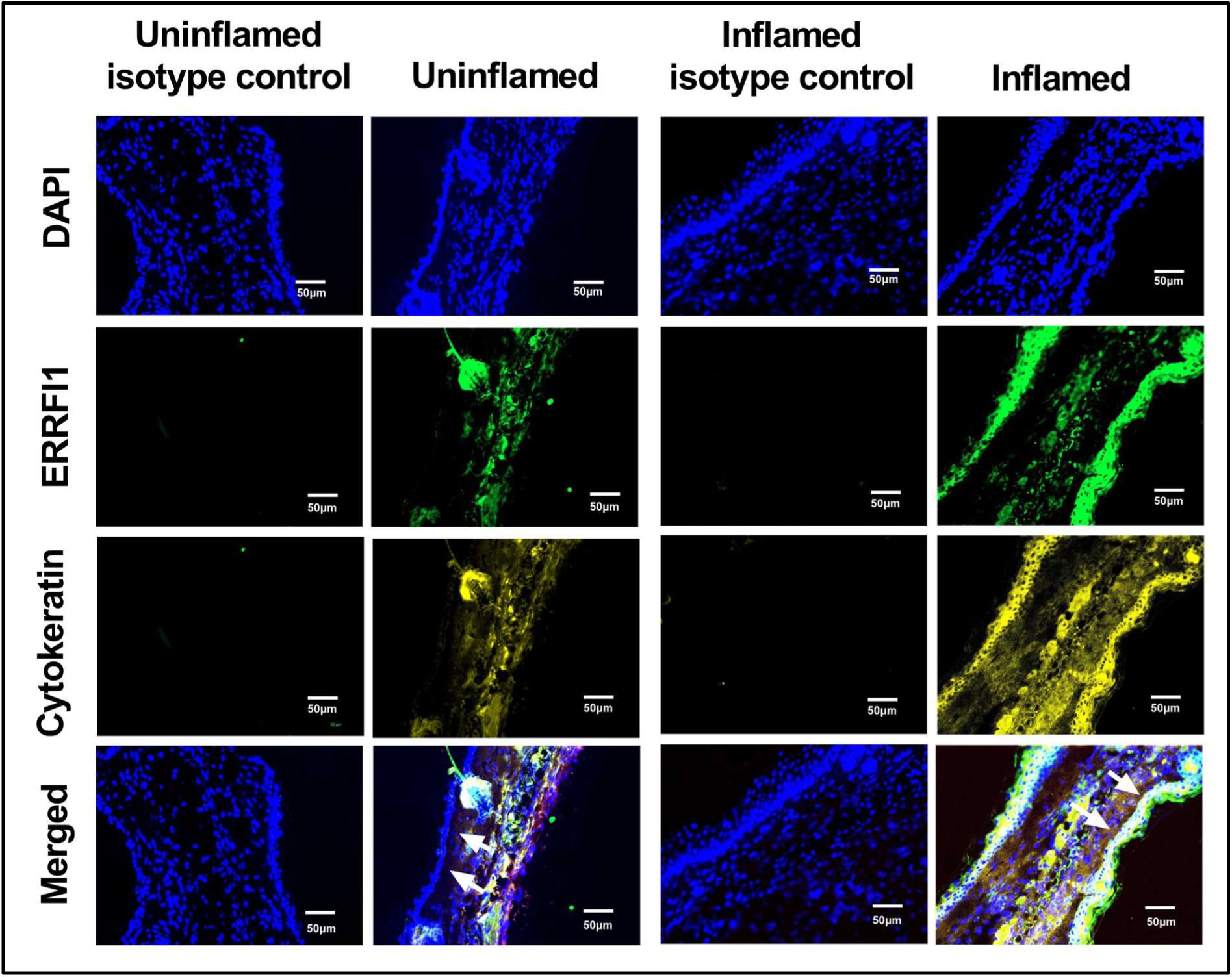
Additional murine ERRFI1 immunofluorescence staining 3.

**Figures S15:**
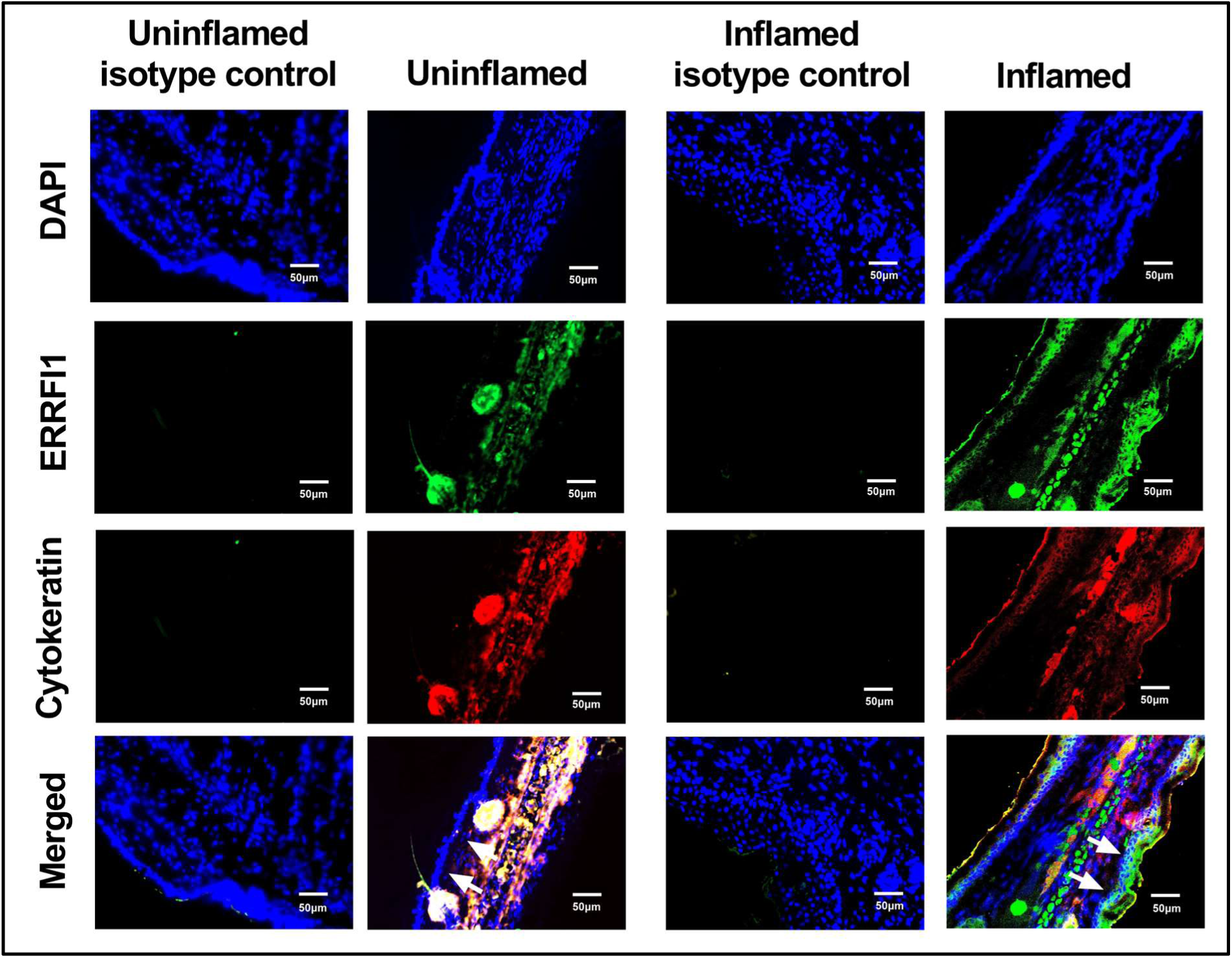
Additional murine ERRFI1 immunofluorescence staining 3.

## Tables

**Table S1:**
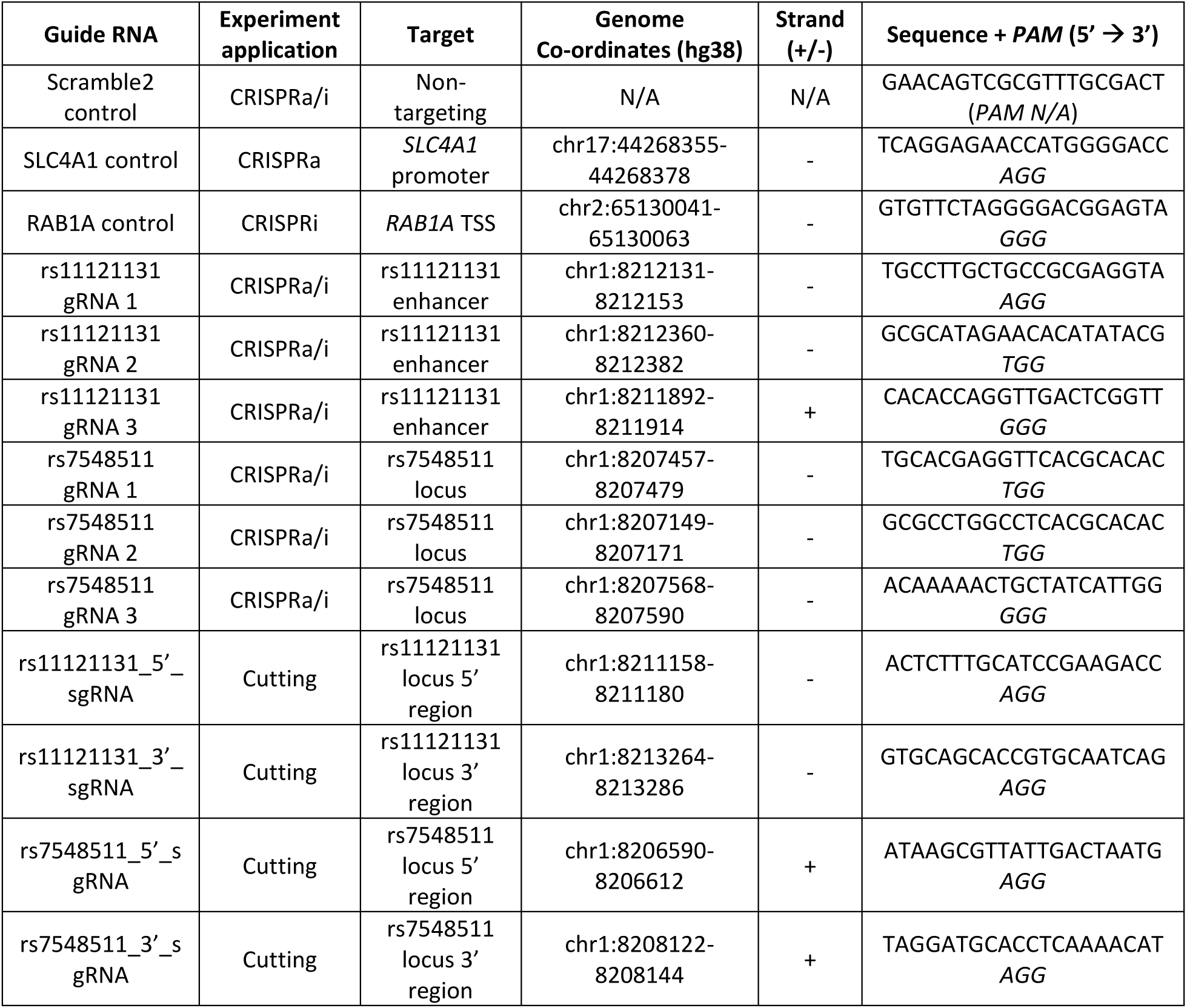
Guide RNAs used in study. Table showing the CRISPR guide RNAs used in this study, along with the associated application (CRISPRa, CRISPRi or cutting), location, strand targeted, and sequence.

**Table S2:**
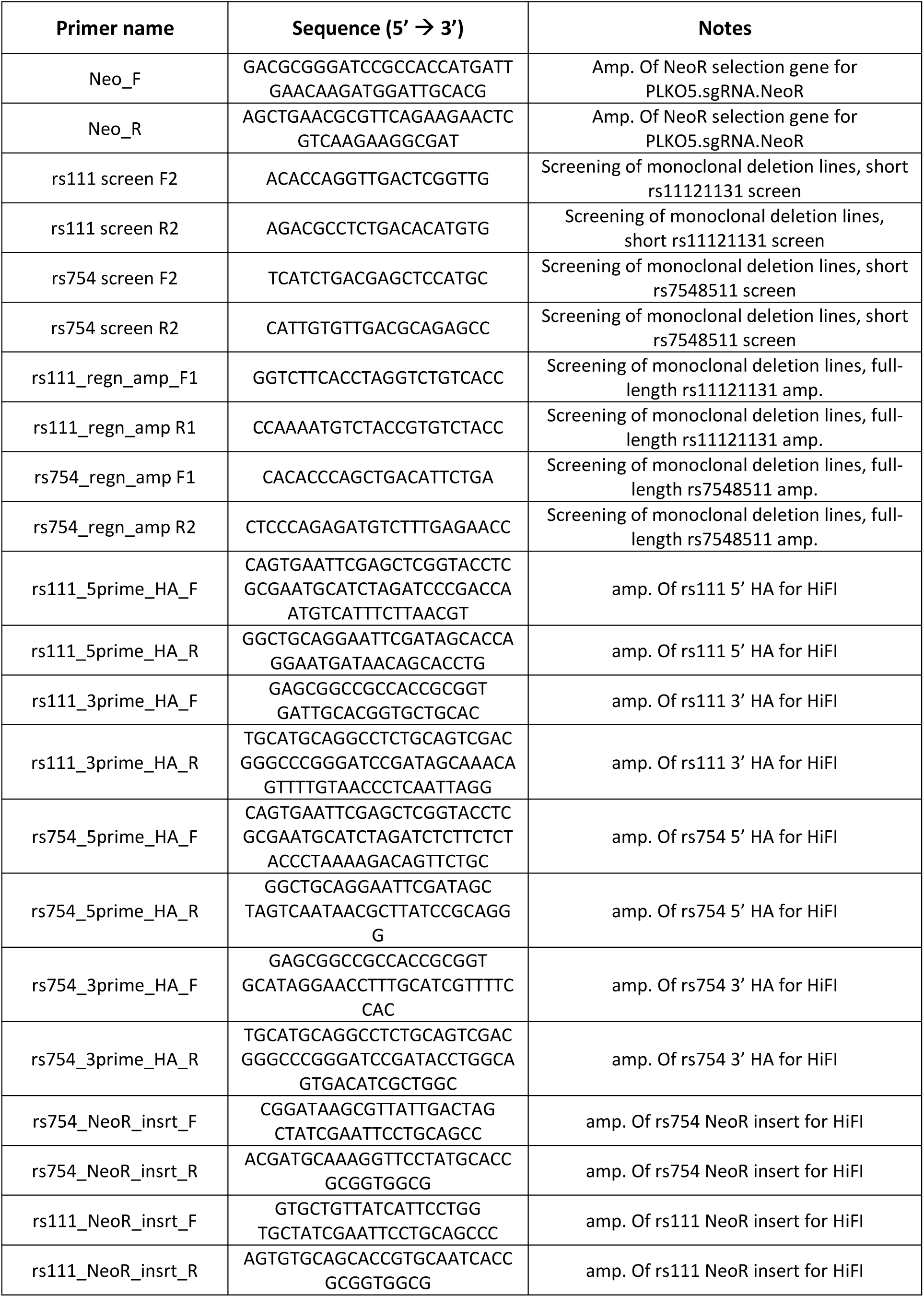
PCR primers used in study.

**Table S3:**
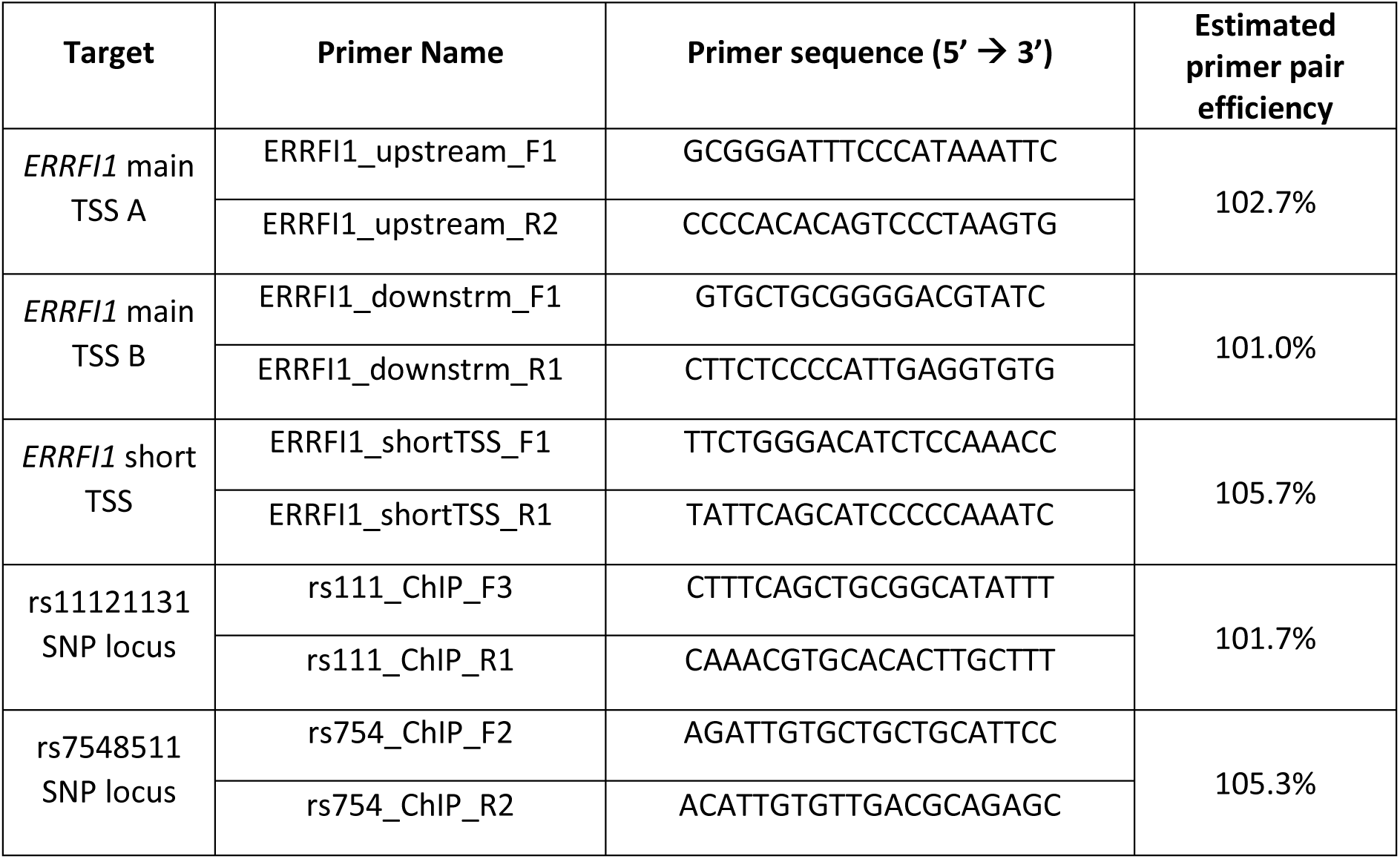
ChIP-qPCR primers.

**Table S4:** TaqMan gene expression assays used in study.

| Gene target | TaqMan Assay ID |
| --- | --- |
| *TBP* | Hs00427620_m1 |
| *YWHAZ* | Hs01122445_g1 |
| *SLC4A1* | Hs00978607_g1 |
| *RAB1A* | Hs00800204_s1 |
| *ERRFI1* | Hs00219060_m1 |
| *TNFRSF9* | Hs00155512_m1 |
| *PARK7* | Hs00994893_g1 |
| *SLC45A1* | Hs00395310_m1 |
| *RERE* | Hs00201558_m1 |
| *EGFR* | Hs01076078_m1 |
| *CCND1* | Hs00765553_m1 |
| *MYC* | Hs00153408_m1 |

**Table S5:** Antibodies used in assessment of murine skin inflammation.

| Panel | Specificity | Conjugate | Clone/cat. no. | Manufacturer |
| --- | --- | --- | --- | --- |
| **Neutrophil stain** | CD45 | BV510 | 30F11 | Biolegend |
|  | CD11b | APC or eF450 | M1/70 | Biolegend |
|  | Gr1 | PECy7 | RB6-8C5 | eBioscience |
| **IL-17 production stain** | CD45 | BV510 | 30F11 | Biolegend |
|  | CD11b | FITC | M1/70 | eBioscience |
|  | CD11c | FITC | N418 | eBioscience |
|  | Gr1 | FITC | RB6-8C5 | eBioscience |
|  | F4/80 | FITC | BM8 | eBioscience |
|  | Ter119 | FITC | TER119 | eBioscience |
|  | FcεRIα | FITC | MAR-1 | eBioscience |
|  | CD19 | FITC | 1D3 | eBioscience |
|  | CD3 | PerCPCy5.5 | 145-2C11 | eBioscience |
|  | TCRγδ | PECy7 | GL3 | eBioscience |
|  | TCRβ | AF700 | H57-597 | Biolegend |
|  | Thy1 | BV786 | 53-2.1 | BD Bioscience |
|  | CD127 | BV711 | SB/199 | BD Bioscience |
|  | IL-17 | PEeF610 | eBio17B7 | eBioscience |

